# Spike generation in electroreceptor afferents introduces additional spectral response components by weakly nonlinear interactions

**DOI:** 10.1101/2024.06.21.599999

**Authors:** Alexandra Barayeu, Maria Schlungbaum, Benjamin Lindner, Jan Grewe, Jan Benda

**Affiliations:** Institute for Neurobiology, Eberhard Karls Universität Tübingen, Auf der Morgenstelle 28, 72076 Tübingen, Germany; Bernstein Center for Computational Neuroscience Berlin, 10115 Berlin, Germany; Department of Physics, Humboldt University Berlin, 12489 Berlin, Germany; Bernstein Center for Computational Neuroscience Tübingen, 72076 Tübingen, Germany

**Keywords:** Volterra series, second-order susceptibility, electric fish, nonlinear coding

## Abstract

Spiking thresholds in neurons or rectification at synapses are essential for neuronal computations rendering neuronal processing inherently nonlinear. Nevertheless, linear response theory has been instrumental for understanding, for example, the impact of noise or neuronal synchrony on signal transmission, or the emergence of oscillatory activity, but is valid only at low stimulus amplitudes or large levels of intrinsic noise. At higher signal-to-noise ratios, however, nonlinear response components become relevant. Theoretical results for leaky integrate-and-fire neurons in the weakly nonlinear regime suggest strong responses at the sum of two input frequencies if one of these frequencies or their sum match the neuron’s baseline firing rate. We here analyze nonlinear responses in two types of primary electroreceptor afferents, the P-units of the active and the ampullary cells of the passive electrosensory system of the wave-type electric fish *Apteronotus leptorhynchus* of either sex. In our combined experimental and modeling approach we identify these predicted nonlinear responses in those 31 out of 172 P-units that are characterized by low intrinsic noise. In contrast, the majority (22 out of 30) ampullary cells show nonlinear responses. Our results provide experimental evidence for nonlinear responses of spike generators in the weakly nonlinear regime. We conclude that such nonlinear responses occur in any sensory neuron that operates in similar regimes particularly at near-threshold stimulus conditions.

**Significance statement:** The generation of action potentials involves a strong threshold nonlinearity. Nevertheless, the encoding of stimuli with small amplitudes by neurons with sufficient intrinsic noise can be well described as a linear system. As the stimulus amplitude is increased, new spectral components start to appear in the so called weakly nonlinear regime. Theory predicts nonlinear interactions whenever one or the sum of two stimulus frequencies matches the neuron’s baseline firing rate. Indeed, we find these interactions in a large set of electrophysiological recordings from primary electroreceptive afferents of a weakly electric fish. The non-linear response components could boost sensory responses to weak signals emitted, for example, by distant conspecifics.

## Introduction

We like to think about signal encoding in terms of linear relations with unique mapping of a given input value to a certain output of the system under consideration. Indeed, such linear methods, for example the transfer function or first-order susceptibility shown in fig. 1 A, have been widely and successfully applied to describe and predict neuronal responses and are an invaluable tools to characterize neural systems (Eggermont et al., 1983; Borst and Theunissen, 1999). Nonlinear mechanisms, on the other hand, are key on different levels of neural processing. Deciding for one action over another is a nonlinear process on the systemic level. On the cellular level, spiking neurons are inherently nonlinear. Whether an action potential is elicited depends on the membrane potential to exceed a threshold (Hodgkin and Huxley, 1952; Koch et al., 1995). Because of such nonlinearities, understanding and predicting neuronal responses to sensory stimuli is in general a difficult task.

**Figure 1.**
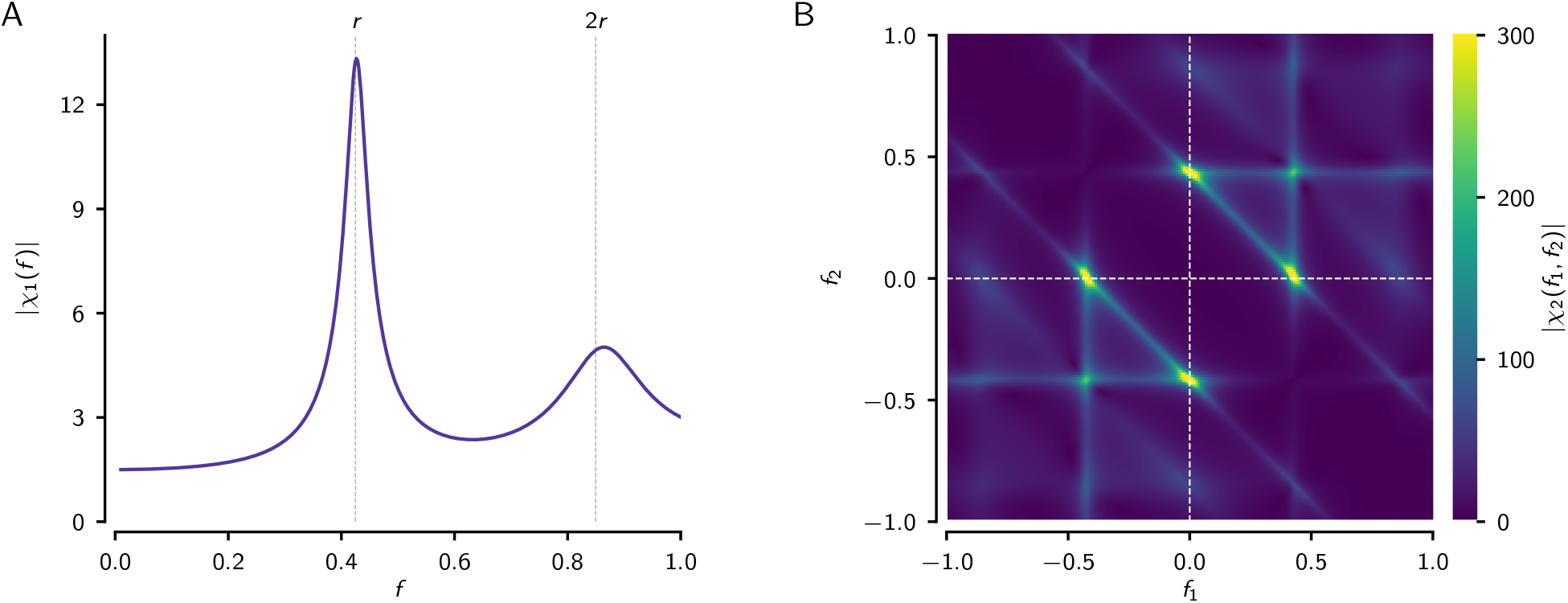
First- and second-order response functions of the leaky integrate-and-fire model. **A** Magnitude of the first-order (linear) response function |*χ*_1_(*f*)|, also known as the “gain” function, quantifies the response amplitude relative to the stimulus amplitude, both measured at the same stimulus frequency. *r* and 2*r* mark the neuron’s baseline firing rate and its harmonic (dashed vertical lines). **B** Magnitude of the second-order (nonlinear) response function |*χ*_2_(*f*_1_, *f*_2_) | quantifies the response at the sum of two stimulus frequencies. Because Fourier spectra have positive and negative frequencies, the sum is actually a difference in the upper-left and lower-right quadrants, since *f*_2_ + *f*_1_ = *f*_2_ −|*f*_1_ |for *f*_1_ *<* 0. For linear systems, the second-order response function is zero, because linear systems do not create new frequencies and thus there is no response at the sum of the two frequencies. The plots show the analytical solutions from Lindner and Schimansky-Geier (2001) and Voronenko and Lindner (2017) with *µ* = 1.1 and *D* = 0.001. Note that the leaky integrate-and-fire model is formulated without dimensions, frequencies are given in multiples of the inverse membrane time constant.

The transfer function that describes the linear properties of a system, is the first-order term of a Volterra series. Higher-order terms successively approximate nonlinear features of a system (Rieke et al., 1999). Second-order kernels have been used in the time domain to predict visual responses in catfish (Marmarelis and Naka, 1972). In the frequency domain, second-order kernels are known as second-order response functions or susceptibilities. Nonlinear interactions of two stimulus frequencies generate peaks in the response spectrum at the sum and the difference of the two. Including higher-order terms of the Volterra series, the nonlinear nature of mammalian visual systems (Victor et al., 1977; Schanze and Eckhorn, 1997), auditory responses in the Torus semicircularis of frogs (Aertsen and Johannesma, 1981), locking in chinchilla auditory nerve fibers (Temchin et al., 2005), spider mechanoreceptors (French et al., 2001), and bursting responses in paddlefish (Neiman and Russell, 2011a) have been demonstrated.

Noise in nonlinear systems, however, linearizes the system’s response properties (Yu and Lewis, 1989; Chialvo et al., 1997). Also, in the limit to small stimuli, nonlinear systems can be well described by linear response theory (Roddey et al., 2000; Doiron et al., 2004; de la Rocha et al., 2007; Sharafi et al., 2013). With increasing stimulus amplitude, the contribution of the second-order kernel of the Volterra series becomes more relevant. For these weakly nonlinear responses analytical expressions for the second-order susceptibility have been derived for leaky-integrate-and-fire (LIF) (Voronenko and Lindner, 2017) and theta model neurons (Franzen et al., 2023). In the suprathreshold regime, in which the LIF generates a baseline firing rate in the absence of an external stimulus, the linear response function has a peak at the baseline firing rate and its harmonics (fig. 1 A) and the second-order susceptibility shows very distinct ridges of elevated nonlinear responses, exactly where one of two stimulus frequencies equals or both frequencies add up to the neuron’s baseline firing rate (fig. 1 B). In experimental data, such structures in the second-order susceptibility have not been reported yet.

Here we search for such weakly nonlinear responses in electroreceptors of the two electrosensory systems of the wave-type electric fish *Apteronotus leptorhynchus*, i.e. the tuberous (active) and the ampullary (passive) electrosensory system. The p-type electroreceptor afferents of the active system (P-units) are driven by the fish’s high-frequency, quasi-sinusoidal electric organ discharges (EOD) and encode disturbances of it (Bastian, 1981). The electroreceptors of the passive system are tuned to lower-frequency exogeneous electric fields such as caused by muscle activity of prey (Kalmijn, 1974). As different animals have different EOD-frequencies, being exposed to stimuli of multiple distinct frequencies is part of the animal’s everyday life (Benda, 2020; Henninger et al., 2020) and weakly nonlinear interactions may occur in the electrosensory periphery. In communication contexts (Walz et al., 2014; Henninger et al., 2018) the EODs of interacting fish superimpose and lead to periodic amplitude modulations (AMs or beats) of the receiver’s EOD. Nonlinear mechanisms in P-units, enable encoding of AMs in their time-dependent firing rates (Bastian, 1981; Walz et al., 2014; Middleton et al., 2006; Barayeu et al., 2023). When multiple animals interact, the superpositions of the EODs induce second-order amplitude modulations referred to as envelopes (Yu et al., 2005; Fotowat et al., 2013; Stamper et al., 2012) and saturation nonlinearities allow also for the encoding of these in the electrosensory periphery (Savard et al., 2011). Field observations have shown that courting males were able to react to the extremely weak signals of distant intruding males despite the strong foreground EOD of the nearby female (Henninger et al., 2018). Weakly nonlinear interactions at particular combinations of signals can be of immediate relevance in such settings as they could boost detectability of the faint signals (Schlungbaum and Lindner, 2023).

## Materials and Methods

### Experimental subjects and procedures

Within this project, we re-evaluated datasets that were recorded between 2010 and 2023 at the Ludwig Maximilian University (LMU) München and the Eberhard-Karls University Tübingen. All experimental protocols complied with national and European law and were approved by the respective Ethics Committees of the Ludwig-Maximilians Universität München (permit no. 55.2-1-54-2531-135-09) and the Eberhard-Karls Unversität Tübingen (permit no. ZP 1/13 and ZP 1/16). The final sample consisted of 172 P-units and 30 ampullary electroreceptor afferents recorded in 80 weakly electric fish of both sexes of the species *Apteronotus leptorhynchus*. Fish were obtained from a commercial supplier for tropical fish (Aquarium Glaser GmbH, Rodgau, Germany) and kept in tanks with a water temperature of 25 ± 1 °C and a conductivity of around 270 µS*/*cm under a 12 h:12 h light-dark cycle.

Before surgery, the animals were deeply anesthetized via bath application of a solution of MS222 (120 mg/l, PharmaQ, Fordingbridge, UK) buffered with Sodium Bicarbonate (120 mg/l). The posterior anterior lateral line nerve (pALLN) was exposed by making a small cut into the skin covering the nerve. The cut was placed dorsal of the operculum just before the nerve descends towards the anterior lateral line ganglion (ALLNG). Those parts of the skin that were to be cut were locally anesthetized by cutaneous application of liquid lidocaine hydrochloride (20 mg/ml, bela-pharm GmbH). During the surgery, water supply was ensured by a mouthpiece to maintain anesthesia with a solution of MS222 (100 mg/l) buffered with Sodium Bicarbonate (100 mg/l). After surgery, fish were immobilized by intramuscular injection of from 25 µl to 50 µl of tubocurarine (5 mg/ml dissolved in fish saline; Sigma-Aldrich). Respiration was then switched to normal tank water and the fish was transferred to the experimental tank.

### Electrophysiological recordings

For the recordings fish were positioned centrally in the experimental tank, with the major parts of their body submerged into the water. Those body parts that were above the water surface were covered with paper tissue to avoid drying of the skin. Local analgesia was refreshed in intervals of two hours by cutaneous application of Lidocaine (2 %; bela-pharm, Vechta, Germany) around the surgical wounds. Electrodes (borosilicate; 1.5 mm outer diameter; GB150F-8P; Science Products, Hofheim, Germany) were pulled to a resistance of 50–100 MΩ (model P-97; Sutter Instrument, Novato, CA) and filled with 1 M KCl solution. Electrodes were fixed in a microdrive (Luigs-Neumann, Ratingen, Germany) and lowered into the nerve. Recordings of electroreceptor afferents were amplified and lowpass filtered at 10 kHz (SEC-05, npi-electronics, Tamm, Germany, operated in bridge mode). All signals, neuronal recordings, recorded EOD, and the generated stimulus, were digitized with sampling rates of 20 or 40 kHz (PCI-6229, National Instruments, Austin, TX). RELACS (https://github.com/relacs/relacs) running on a Linux computer was used for online spike and EOD detection, stimulus generation, and calibration. Recorded data was then stored on the hard drive for offline analysis.

### Identification of P-units and ampullary cells

Recordings were classified as P-units if baseline action potentials phase locked to the EOD with vectors strengths between 0.7 and 0.95, a baseline firing rate larger than 30 Hz, a serial correlation of subsequent interspike intervals below zero, a coefficient of variation of baseline interspike intervals below 1.5 and during stimulation below 2. P-units are clearly distinguished from T-type electroreceptors, that we did not analyze here, by having firing rates much lower than the EOD frequency of the fish (no 1:1 locking to the EOD). As ampullary cells we classified recordings with vector strengths below 0.15, baseline firing rate above 10 Hz, baseline CV below 0.18, CV during stimulation below 1.0, and a response modulation during stimulation below 80 Hz (Grewe et al., 2017). We here selected only those cells of which the neuron’s baseline activity as well as the responses to band-limited white noise stimuli were recorded.

### Electric field recordings

For monitoring the EOD without the stimulus, two vertical carbon rods (11 cm long, 8 mm diameter) in a head-tail configuration were placed isopotential to the stimulus. Their signal was differentially amplified with a gain factor between 100 and 500 (depending on the recorded animal) and band-pass filtered (3 to 1500 Hz pass-band, DPA2-FX; npi electronics, Tamm, Germany). For an estimate of the transdermal potential that drives the electroreceptors, two silver wires spaced by 1 cm were located next to the left gill of the fish and orthogonal to the fish’s longitudinal body axis (amplification 100 to 500 times, band-pass filtered with 3 to 1 500 Hz pass-band, DPA2-FX; npi-electronics, Tamm, Germany). This local EOD measurement recorded the combination of the fish’s own EOD and the applied stimulus.

### Stimulation

Electric stimuli were attenuated (ATN-01M, npi-electronics, Tamm, Germany), isolated from ground (ISO-02V, npi-electronics, Tamm, Germany) and delivered via two horizontal carbon rods (30 cm length, 8 mm diameter) located 15 cm parallel to each side of the fish. The fish were stimulated with band-limited Gaussian white noise stimuli, i.e. signals with equal power at all frequencies up to a cut-off frequency and with a stationary Gaussian probability density. For the ampullary cells we chose a cut-off frequency of 150 Hz, whereas for the P-units we used either 300 or 400 Hz. The stimuli were generated by drawing normally distributed real and imaginary numbers for all frequencies up to the desired cut-off frequency in the Fourier domain and then applying an inverse Fourier transform (Billah and Shinozuka, 1990; Škorjanc et al., 2023). The stimulus intensity is given as a contrast, i.e. the standard deviation of the resulting amplitude modulation relative to the fish’s EOD amplitude. The contrast varied between 1 and 20 % (median 5 %) for P-units and 2.5 and 20 % (median 5 %) for ampullary cells. Only recordings with noise stimuli with a duration of at least 2 s (maximum of 50 s, median 10 s) and enough repetitions to results in at least 100 FFT segments (see below, P-units: 100–1520, median 313, ampullary cells: 105 – 3648, median 722) were included into the analysis. When ampullary cells were recorded, the noise stimuli *s*(*t*) were directly applied as the stimulus and thus were simply added to the fish’s own EOD: *s*(*t*) + *EOD*(*t*). To create random amplitude modulations (RAM) for P-unit recordings, the noise stimulus was first multiplied with the EOD of the fish (MXS-01M; npi electronics) and then played back through the stimulation electrodes: *EOD*(*t*) + *s*(*t*)*EOD*(*t*) = (1 + *s*(*t*))*EOD*(*t*).

## Data analysis

Data analysis was done in Python 3 using the packages matplotlib (Hunter, 2007), numpy (Walt et al., 2011), scipy (Virtanen et al., 2020), nixio (Stoewer et al., 2014), and thunderlab (https://github.com/bendalab/thunderlab).

### Code accessibility

The P-unit model parameters and spectral analysis algorithms are available at https://github.com/bendalab/punitmodel/tree/v1.

### Baseline analysis

The baseline firing rate *r* was calculated as the number of spikes divided by the duration of the baseline recording (median 32 s). The coefficient of variation (CV) of the interspike intervals (ISI) is their standard deviation relative to their mean: 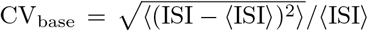. If the baseline was recorded several times in a recording, the measures from the longest recording were taken.

### White noise analysis

When stimulated with band-limited white noise stimuli, neuronal activity is modulated around the average firing rate that is similar to the baseline firing rate and in that way encodes the time-course of the stimulus. For an estimate of the time-dependent firing rate *r*(*t*) we convolved each spike train with normalized Gaussian kernels with standard deviation of 1 ms and averaged the resulting single-trail firing rates over trials. The response modulation quantifies the variation of *r*(*t*) computed as the standard deviation in time 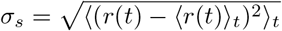, where ⟨ · ⟩_*t*_ denotes averaging over time.

### Spectral analysis

To characterize the relation between the spiking response evoked by white-noise stimuli, we estimated the first- and second-order susceptibilities in the frequency domain. For this we converted spike times into binary vectors *x*_*k*_ with Δ*t* = 0.5 ms wide bins that are set to 2 kHz where a spike occurred and zero otherwise. Fast Fourier transforms (FFT) *S*(*ω*) and *X*(*ω*) of the stimulus *s*_*k*_ (also down-sampled to a sampling rate of 2 kHz) and *x*_*k*_, respectively, were computed numerically according to

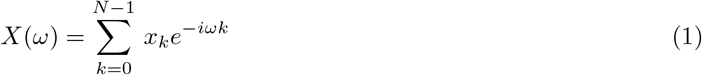

for *N* = 512 long segments of *T* = *N* Δ*t* = 256 ms duration with no overlap, resulting in a spectral resolution of about 4 Hz. In fig. 4 we used *N* = 2^18^ and Δ*t* = 0.1 ms (26.2 s). Note, that for a real Fourier integral a factor Δ*t* is missing. For simplicity we use angular frequencies *ω* = 2*πf* instead of frequencies *f*. Also note, that the Fourier transform of a signal results in both positive as well as negative frequencies. This is because the signal is decomposed into complex exponentials, not sine or cosine waves. To get back a real-valued sine or cosine wave, the components at positive and negative frequencies need to be combined. For example, the sum *e*^*−ωt*^ + *e*^+*ωt*^ of two complex exponentials at frequencies ± *ω* is 2 cos(*ωt*).

In the experimental data the duration of the noise stimuli varied and they were presented once or repeatedly (frozen noise). For the analysis we discarded the responses within the initial 200 ms of stimulation in each trial. To make the recordings comparable we always used the first 100 segments from as many trials as needed for the following analysis.

In the simulations we generated for each trial a new realization of the noise stimulus. We discarded the first 500 ms of the response and used the following 10 FFT segments for the analysis. This was repeated until 10^6^ or 10^7^ FFT segments were collected.

The power spectrum of the stimulus *s*(*t*) was estimated as

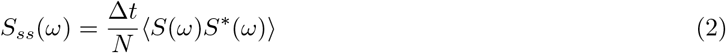

with *S*^*∗*^ being the complex conjugate of *S* and ⟨ · ⟩ denoting the average over the FFT segments. The factor in front is Δ*t*^2^ from the missing integration factors of the two Fourier transforms, Eq. (1), divided by *T* = *N* Δ*t*, needed to make this a proper power spectral density.

The power spectrum of the spike trains *S*_*xx*_(*ω*) was estimated accordingly. Likewise, the cross-spectrum *S*_*xs*_(*ω*) between stimulus and evoked spike trains was estimated according to

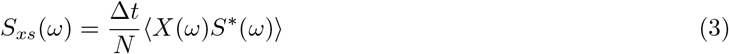

The first-order susceptibility (transfer function)

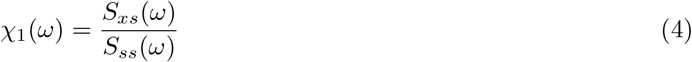

was then computed from *S*_*xs*_(*ω*) and *S*_*ss*_(*ω*). We report *χ*_1_(*ω*) in Hz/%, i.e. firing rate per percent stimulus contrast. Multiplying *χ*_1_(*ω*) with the contrast of a sinusoidal stimulus in percent results in the amplitude of the evoked firing rate modulation in Hertz.

The second-order cross-spectrum

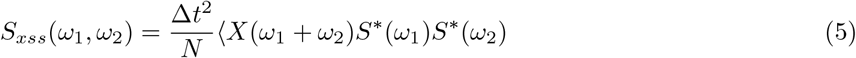

quantifies nonlinear interactions that generate responses at the sum and difference (for negative *ω*_1_ or *ω*_2_) evoked by two stimulus frequencies *ω*_1_ and *ω*_2_. The second-order susceptibility

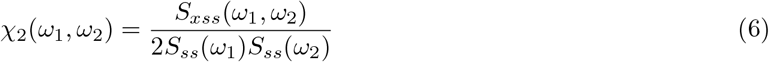

normalizes the second-order cross-spectrum by the spectral power at the two stimulus frequencies. We report *χ*_2_(*ω*_1_, *ω*_2_) in Hz/%^2^, i.e. firing rate per percent stimulus contrast squared. Throughout the manuscript we only show the absolute values of the complex-valued second-order susceptibility matrix and ignore the corresponding phases.

### Susceptibility index

We expected to see a sharp ridge in the second-order susceptibility at *ω*_1_ + *ω*_2_ = *r* (Voronenko and Lindner, 2017; Franzen et al., 2023). To characterize this in a single number we computed a susceptibility index. First, we projected the absolute values of the second-order susceptibility matrix onto the diagonal by averaging over anti-diagonal elements. In this projection *D*(*f*) we took the position of the maximum

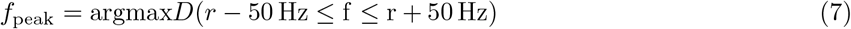

within ± 50 Hz of the neuron’s baseline firing rate *r* as the position of the expected peak. For an estimate of the noise-floor surrounding this peak we averaged over 10 Hz wide windows 10 Hz to the left and right of the peak:

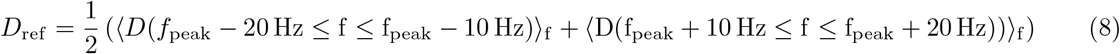

The size of the peak relative to this reference is then the susceptibility index

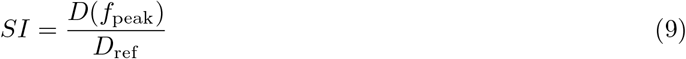

Values larger than one indicate a sharp ridge in the susceptibility matrix close to where the stimulus frequencies add up to the baseline firing rate.

### Leaky integrate-and-fire models for P-units

Modified leaky integrate-and-fire (LIF) models were constructed to reproduce the specific firing properties of P-units (Chacron et al., 2001; Sinz et al., 2020; Barayeu et al., 2023). Its basic components (static nonlinearity, low-pass filtering and spike generation) are equivalent to models of hair cells in auditory systems (Eggermont et al., 1983). The sole driving input into the P-unit model during baseline, i.e. when no external stimulus was given, is the fish’s own EOD, modeled as a cosine wave

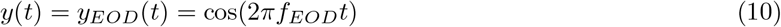

with EOD frequency *f*_*EOD*_ and an amplitude of one.

To mimic the interaction with other fish, the EODs of a second or third fish with EOD frequencies *f*_1_ and *f*_2_, respectively, were added to the normalized EOD, Eq. (10), of the receiving fish according to their contrasts, *c*_1_ and *c*_2_ at the position of the receiving fish:

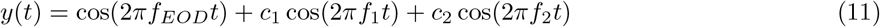

For two fish, *c*_2_ = 0.

Random amplitude modulations (RAMs) were simulated by first generating the AM as a band-limited white noise stimulus *s*(*t*). For this, random real and imaginary numbers were drawn from normal distributions for each frequency component in the range from 0 to 300 Hz in the Fourier domain (Billah and Shinozuka, 1990; Škorjanc et al., 2023). By means of the inverse Fourier transform, the time course of the RAM stimulus, *s*(*t*), was generated. The input to the model was then

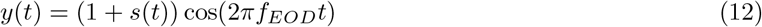

The contrast *c* of the RAM is the standard deviation of the RAM relative to the amplitude of the receiving fish.First, the input *y*(*t*) is thresholded by setting negative values to zero:

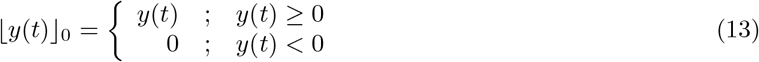

(fig. 2 B). This threshold models the transfer function of the synapses between the primary receptor cells and the afferent (Moser and Starr, 2016) and may also include nonlinear properties introduced by low-threshold Kv1 channels known to be present in the afference (Smith et al., 2006; Nogueira and Caputi, 2013). Together with a low-pass filter

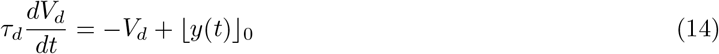

the threshold operation is required for extracting the amplitude modulation from the input (Barayeu et al., 2024). As detailed in the discussion, the intricacies of the threshold nonlinearity Eq. (13) and the low-pass filter Eq. (14) do not matter here, as all cross-spectra were computed between the spiking response and the amplitude-modulation *s*(*t*) — and not the full input signal *y*(*t*), Eq. (12).

**Figure 2.**
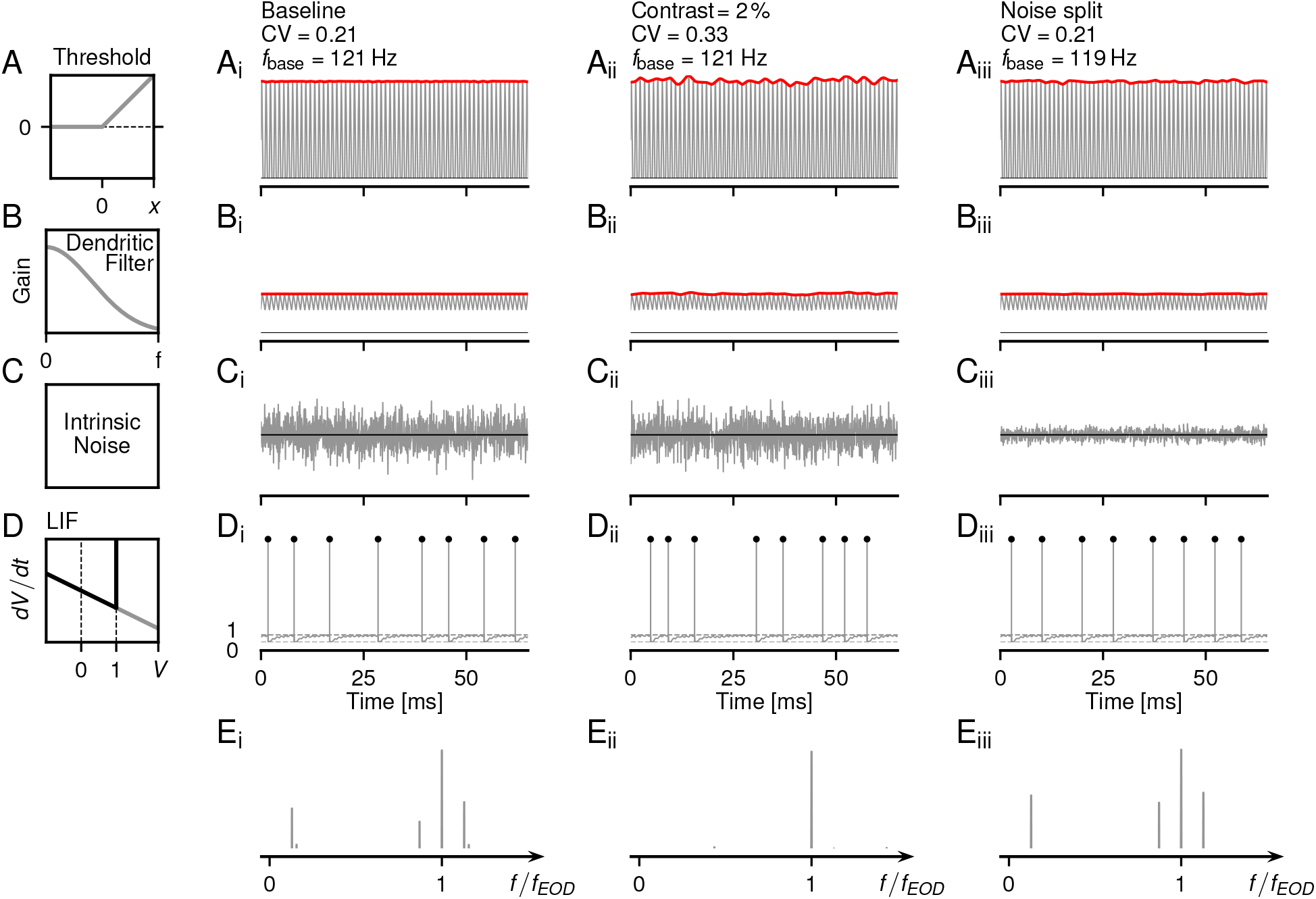
Architecture of the P-unit model. Each row illustrates subsequent processing steps for three different stimulation regimes: (i) baseline activity without external stimulus, only the fish’s self-generated EOD (the carrier, Eq. (10)) is present. (ii) RAM stimulation, Eq. (12). The amplitude of the EOD carrier is modulated with a weak (2 % contrast) band-limited white-noise stimulus. (iii) Noise split, Eqs. (18)–(20), where 90 % of the intrinsic noise is replaced by a RAM stimulus, whose amplitude is scaled to maintain the mean firing rate and the CV of the ISIs of the model’s baseline activity. As an example, simulations of the model for cell “2012-07-03-ak” are shown. **A** The stimuli are thresholded, Eq. (13), by setting all negative values to zero. **B** Subsequent low-pass filtering, Eq. (14), attenuates the carrier and carves out the AM signal. **C** Intrinsic Gaussian white-noise is added to the signals shown in B. Note the reduced internal noise amplitude in the noise split (iii) condition. **D** Spiking output of the LIF model, Eqs. (15)–(17), in response to the sum of B and C. **E** Power spectra of the LIF neuron’s spiking activity. Both, baseline activity (E_i_) and noise split (E_iii_), have the same peaks in the response spectrum at *r, f*_*EOD*_ −*r, f*_*EOD*_, and *f*_*EOD*_ + *r*. With RAM stimulation (E_ii_), the peak at the baseline firing rate, *r*, is washed out.

The low-pass filter models passive signal conduction in the afferent’s dendrite (fig. 2 B) from the synapse to the spike initiation zone. *τ*_*d*_ is the effective membrane time constant of this part of the dendrite. Dendritic low-pass filtering was also necessary to reproduce the loose coupling of P-unit spikes to the EOD while maintaining high sensitivity at small amplitude modulations.

The dendritic voltage *V*_*d*_(*t*) is then fed into a stochastic leaky integrate-and-fire (LIF) model with adaptation,

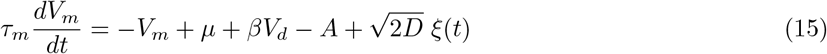

where *τ*_*m*_ is the membrane time-constant, *µ* is a fixed bias current, *β* is a scaling factor for *V*_*d*_, *A* is an inhibiting adaptation current, and 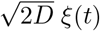 is a Gaussian white noise with strength *D*. Note, that all state variables, membrane voltages *V*_*d*_ and *V*_*m*_ as well as the adaptation current *A*, are dimensionless.

The adaptation current *A* follows

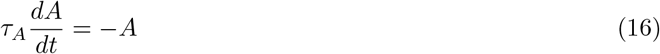

with adaptation time constant *τ*_*A*_ (Benda and Herz, 2003; Benda et al., 2005, 2010).

Whenever the membrane voltage *V*_*m*_(*t*) crosses the spiking threshold *θ* = 1, a spike was generated, *V*_*m*_(*t*) was reset to 0, the adaptation current was incremented by Δ*A/τ*_*A*_, and integration of *V*_*m*_(*t*) was paused for the duration of a refractory period *t*_*ref*_ (fig. 2 D):

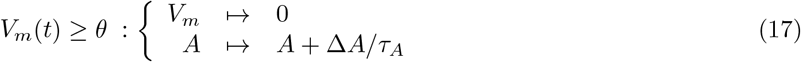

The P-unit models were integrated by the Euler forward method with a time-step of Δ*t* = 0.05 ms. For each trial of a simulation, *V*_*m*_ was drawn from a uniform distribution between 0 and 1 and the initial value of *A* was jittered by adding a random number drawn from a normal distribution with standard deviation of 2 % of its initial value. Then the first 500 ms of any simulation were discarded to remove remaining transients.

The eight free parameters of the P-unit model *β, τ*_*m*_, *µ, D, τ*_*A*_, Δ_*A*_, *τ*_*d*_, and *t*_*ref*_, were fitted to both the baseline activity (baseline firing rate, CV of ISIs, serial correlation of ISIs at lag one, and vector strength of spike coupling to EOD) and the responses to step increases and decreases in EOD amplitude (onset and steady-state responses, effective adaptation time constant, Benda et al., 2005) of recorded P-units. Model parameters of all 39 cells are summarized in file models.csv of our punitmodel repository at https://github.com/bendalab/punitmodel/tree/v1.

### Noise split

Based on the Furutsu-Novikov theorem (Furutsu, 1963; Novikov, 1965; Lindner, 2022; Egerland, 2021), we split the total noise, 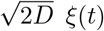, of a LIF model, Eq. (15), into two parts. The first part is the intrinsic noise term,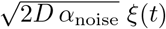, whose strength is reduced by a factor *α*_noise_ = 0.1 (fig. 2 C_iii_). The second part replaces the now missing intrinsic noise by a driving input signal *s*_*ξ*_(*t*), a RAM stimulus with frequencies up to 300 Hz (fig. 2 A_iii_). The LIF model with noise split then reads

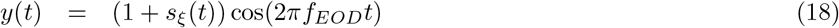

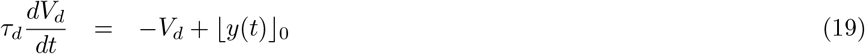

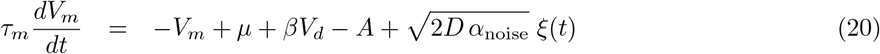

Both, the reduced intrinsic noise and the RAM stimulus, need to replace the original intrinsic noise. Because the RAM stimulus is band-limited and undergoes some transformations before it is added to the reduced intrinsic noise, it is not *a priori* clear, what the amplitude of the RAM should be. We bisected the amplitude of *s*_*ξ*_(*t*), until the CV of the resulting interspike intervals matched the one of the original model’s baseline activity. The second-order cross-spectra, Eq. (5), were computed between the RAM stimulus *s*_*ξ*_(*t*) and the spike train *x*(*t*) it evoked. In this way, the effective signal-to-noise ratio can be increased while maintaining the total noise in the system.

## Results

We explored a large set of electrophysiological data from primary afferents of the active and passive electrosensory system, P-units and ampullary cells (Grewe et al., 2017; Hladnik and Grewe, 2023), that were recorded in the brown ghost knifefish *Apteronotus leptorhynchus*. We re-analyzed this dataset to search for weakly nonlinear responses that have been predicted in previous theoretical work (Voronenko and Lindner, 2017). Additional simulations of LIF-based models of P-unit spiking help to interpret the experimental findings in this theoretical framework. We start with demonstrating the basic concepts using example P-units and respective models and then compare the population of recordings in both cell types.

### Nonlinear responses in P-units stimulated with two frequencies

Without external stimulation, a P-unit is driven by the fish’s own EOD alone (with a specific EOD frequency *f*_EOD_) and spontaneously fires action potentials at the baseline rate *r*. Accordingly, the power spectrum of the baseline activity has a peak at *r* (fig. 3 A). In the communication context, this animal (the receiver) is exposed to the EODs of one or many foreign fish. Superposition of the receiver’s EOD with an EOD of another fish with frequency *f*_1_ results in a beat, a periodic amplitude modulation of the receiver’s EOD. The frequency of the beat is given by the difference frequency Δ*f*_1_ = *f*_1_ − *f*_EOD_ between the two fish. P-units encode this beat in their firing rate (Bastian, 1981) and consequently the power spectrum of the response has a peak at the beat frequency (fig. 3 B). A second peak at the first harmonic of the beat frequency is indicative of a nonlinear process that here is associated with the clipping of the P-unit’s firing rate at zero (Barayeu et al., 2023). Pairing the fish with another fish at a higher beat frequency Δ*f*_2_ = *f*_2_ −*f*_EOD_ *>* Δ*f*_1_ results in a weaker response with a single peak in the response power spectrum, suggesting a linear response (fig. 3 C). The weaker response to this beat can be explained by the beat tuning of the cell (Walz et al., 2014). Note, Δ*f*_2_ has been deliberately chosen to match the recorded P-unit’s baseline firing rate.

**Figure 3.**
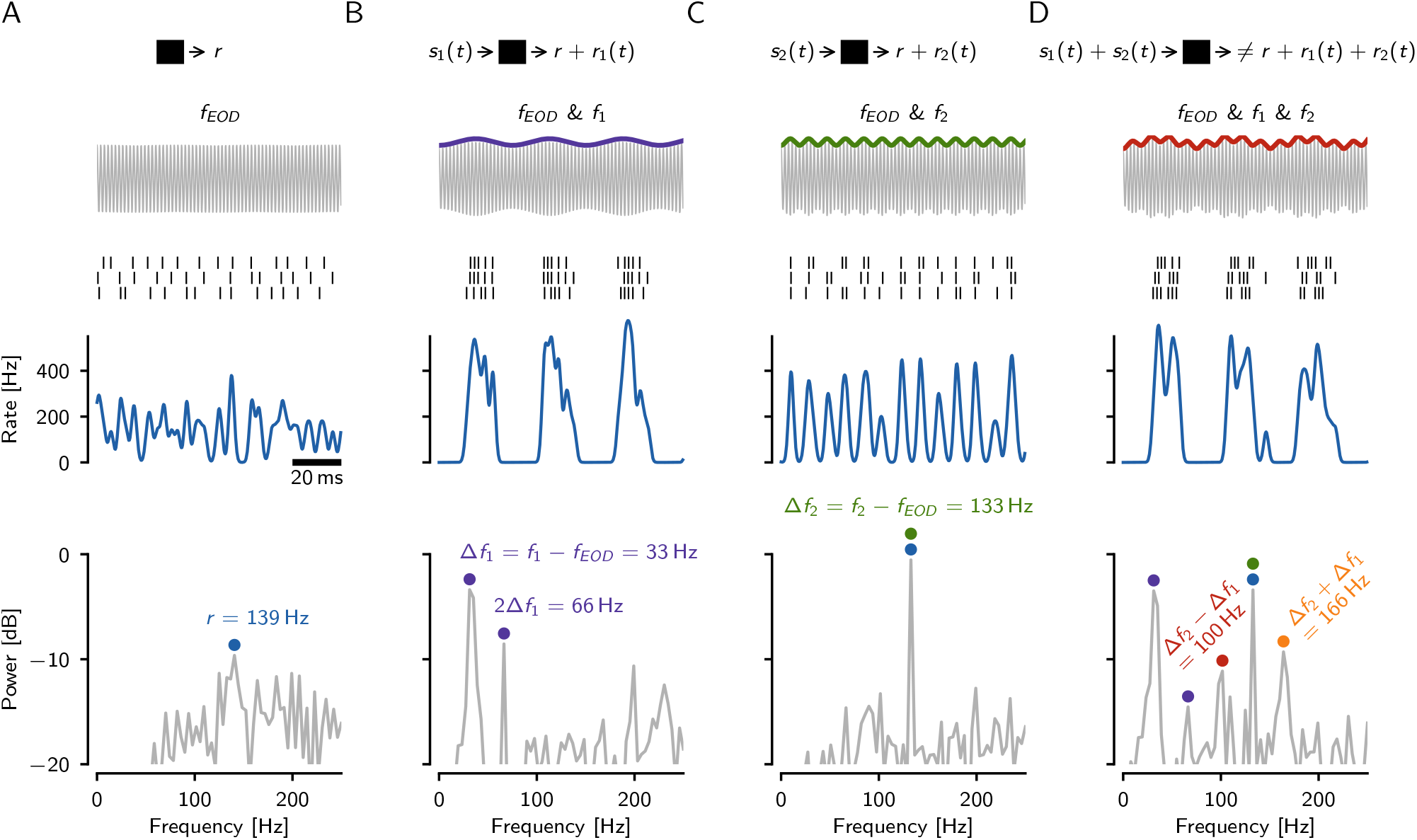
Nonlinearity in an electrophysiologically recorded P-unit of *A. leptorhynchus* in a three-fish setting (cell identifier “2021-08-03-ac”). Receiver with EOD frequency *f*_EOD_ = 664 Hz encounters fish with EOD frequencies *f*_1_ = 631 Hz and *f*_2_ = 797 Hz. Both foreign signals have the same strength relative to the own field amplitude (10 % contrast). Top row: Sketch of signal processing in the nonlinear system (black box). Second row: Superposition of the receiver EOD with the EODs of other fish, colored line highlights the amplitude modulation. Third row: Three trials of spike trains of the recorded P-unit. Fourth row: Firing rate, estimated by convolution of the spike trains with a Gaussian kernel. Bottom row: Power spectrum of the spike trains. **A** Baseline condition: The cell is driven by the self-generated field alone. The baseline firing rate *r* = 139 Hz dominates the power spectrum (blue circle). **B** The receiver’s EOD and a single conspecific with an EOD frequency *f*_1_ = 631 Hz are present. Superposition of the two EODs induces a periodic amplitude modulation, referred to as beat, with beat frequency Δ*f*_1_ = 33 Hz. The P-unit strongly responds to this beat (purple). **C** The receiver and a fish with an EOD frequency *f*_2_ = 797 Hz are present. The resulting beat Δ*f*_2_ = 133 Hz is faster as the difference between the EOD frequencies is larger. The P-unit response to this faster beat is weaker (green). **D** All three fish with EOD frequencies *f*_EOD_, *f*_1_, and *f*_2_ are present. Additional peaks occur in the power spectrum of the spike response at the sum (orange) and difference (red) of the two beat frequencies, indicating nonlinear interactions between the two frequencies in the P-unit. Note, the spectrum of the raw signal (top row, gray) has power only at the three EOD frequencies *f*_*EOD*_, *f*_1_, and *f*_2_.

When stimulating with both foreign signals simultaneously, additional peaks appear in the response power spectrum at the sum Δ*f*_1_ +Δ*f*_2_ and the difference frequency Δ*f*_2_ − Δ*f*_1_ (fig. 3 B). Thus, the cellular response is not equal to the sum of the responses to the two beats presented separately. These additional peaks at the sum and the difference of the two stimulus frequencies are a hallmark of nonlinear interactions that, by definition, are absent in linear systems.

### Linear and weakly nonlinear regimes

The stimuli used in fig. 3 had the same AM contrasts of 10 % that evoke rather strong modulations in a P-unit’s firing rate response. Whether these definitely not-small stimulus amplitudes fall into the weakly nonlinear regime as in Voronenko and Lindner (2017) is not clear. In order to illustrate how the responses to two beat frequencies develop over a range of amplitudes we use a stochastic leaky-integrate-and-fire (LIF) based P-unit model fitted to a specific electrophysiologically measured cell (Barayeu et al., 2023).

At very low stimulus contrasts (in the example cell less than approximately 1.2 % relative to the receiver’s EOD amplitude) the spectrum has small peaks at the beat frequencies (fig. 4 A,B, green and purple). The amplitudes of these peaks initially increase linearly with stimulus amplitude (fig. 4 E, thin lines), an indication of the linear response at lowest stimulus amplitudes. The largest peak at the receiver’s EOD frequency indicates the stochastic locking of the P-unit spikes to the EOD, and its side-peaks are remnants of the two stimulating EOD frequencies (Sinz et al., 2020).

**Figure 4.**
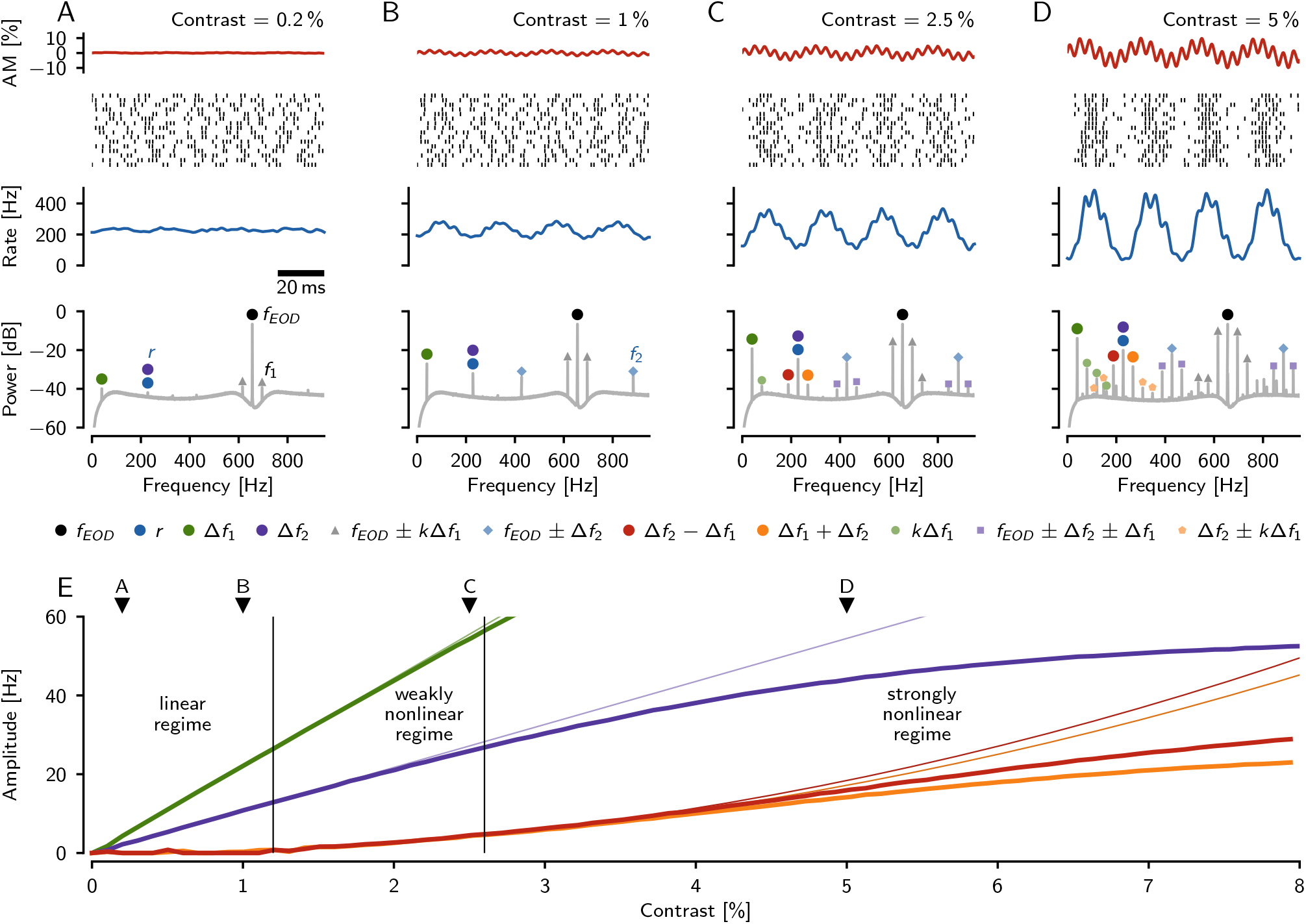
Linear and nonlinear responses of a model P-unit in a three-fish setting in dependence on stimulus amplitudes. The model P-unit (identifier “2018-05-08-ad”) was stimulated with two sine waves of equal amplitude (contrast) at difference frequencies Δ*f*_1_ = 40 Hz and Δ*f*_2_ = 228 Hz relative the receiver’s EOD frequency *f*_*EOD*_ = 656 Hz. Δ*f*_2_ was set to match the baseline firing rate *r* of the P-unit. **A–D** Top row: the stimulus, an amplitude modulation of the receiver’s EOD resulting from the stimulation with the two sine waves. The contrasts of both beats increase from A to D as indicated. Second row: Spike raster of the model P-unit response. Third row: average firing rate and, bottom row: power spectrum estimated from 1000 trials of spike train responses. **A** At low stimulus contrasts the response to the two beat frequencies is linear. These frequencies are present in the response spectrum (green and purple circles), the peak at Δ*f*_2_ (purple) enhances the peak at baseline firing rate (blue). The largest peak, however, is always the one at the EOD frequency of the receiver (black), reflecting the locking of P-unit spikes to the receiver’s own EOD. The peak at *f*_*EOD*_ is also flanked by harmonics of Δ*f*_1_ (gray triangles), where *f*_*EOD*_ + Δ*f*_1_ = *f*_1_ is the EOD frequency of the first fish. **B** At moderately higher stimulus contrast, the peaks in the response spectrum at the two beat frequencies become larger. In addition, two peaks at *f*_*EOD*_ − Δ*f*_2_ and *f*_*EOD*_ +Δ*f*_2_ appear (blue diamonds), the latter is the EOD frequency of the second fish, *f*_2_. These peaks indicate nonlinear processes acting on the three EOD frequencies from which the beat frequencies are generated. **C** At intermediate stimulus contrasts, nonlinear responses start to appear at the sum and the difference of the beat frequencies (orange and red circles). Similarly, side peaks appear at± Δ*f*_1_ around *f*_*EOD*_± Δ*f*_2_ (purple squares) as well as harmonics of Δ*f*_1_ (small green circles). **D** At higher stimulus contrasts many more peaks appear in the power spectrum. Most of them are interactions of *f*_*EOD*_ and Δ*f*_2_ with harmonics of Δ*f*_1_ (*k* = 1, 2, *…*), since the cell responds strongest to Δ*f*_1_. **E** Amplitude of the linear (at Δ*f*_1_ and Δ*f*_2_) and nonlinear (at Δ*f*_2_ − Δ*f*_1_ and Δ*f*_1_ + Δ*f*_2_) responses of the model P-unit as a function of beat contrast (thick lines). Thin lines indicate the initial linear and quadratic dependence on stimulus amplitude for the linear and nonlinear responses, respectively. In the linear regime, below a stimulus contrast of about 1.2 % (left vertical line), the only peaks in the response spectrum are at the stimulus frequencies. In the weakly nonlinear regime up to a contrast of about 3.5 % peaks arise at the sum and the difference of the two stimulus frequencies. At stronger stimulation the amplitudes of these nonlinear responses deviate from the quadratic dependency on stimulus contrast.

This linear regime is followed by the weakly nonlinear regime (in the example cell between approximately 1.2 % and 2.6 % stimulus contrast). In addition to the peaks at the stimulating beat frequencies, peaks at the sum and the difference of the beat frequencies appear in the response spectrum (fig. 4 C, orange and red). The amplitudes of these two peaks initially increase quadratically with stimulus amplitude (fig. 4 E, thin lines). Note, that we have chosen Δ*f*_2_ to match the baseline firing rate *r* of the neuron.

At higher stimulus amplitudes, the linear response and the weakly-nonlinear response begin to deviate from their linear and quadratic dependency on amplitude (fig. 4 E) and additional peaks appear in the response spectrum (fig. 4 D). At high stimulus contrasts, additional nonlinearities in the system, in particular clipping of the firing rate, shape the responses.

For this example, we chose very specific stimulus (beat) frequencies. In the following, however, we are interested in how the nonlinear responses depend on different combinations of stimulus frequencies in the weakly nonlinear regime. For the sake of simplicity we will drop the Δ notation even though P-unit stimuli are beats.

### Nonlinear signal transmission in P-units

P-units fire action potentials probabilistically phase-locked to the self-generated EOD (Bastian, 1981). Skipping of EOD cycles leads to the characteristic multimodal ISI distribution with maxima at integer multiples of the EOD period (fig. 5 A). In this example, the baseline ISI distribution has a CV_base_ of 0.49, which is at the center of the P-unit population (Hladnik and Grewe, 2023). Spectral analysis of the baseline activity shows two major peaks: the first is located at the baseline firing rate *r*, the second is located at the discharge frequency *f*_EOD_ of the electric organ (fig. 5 B).

**Figure 5.**
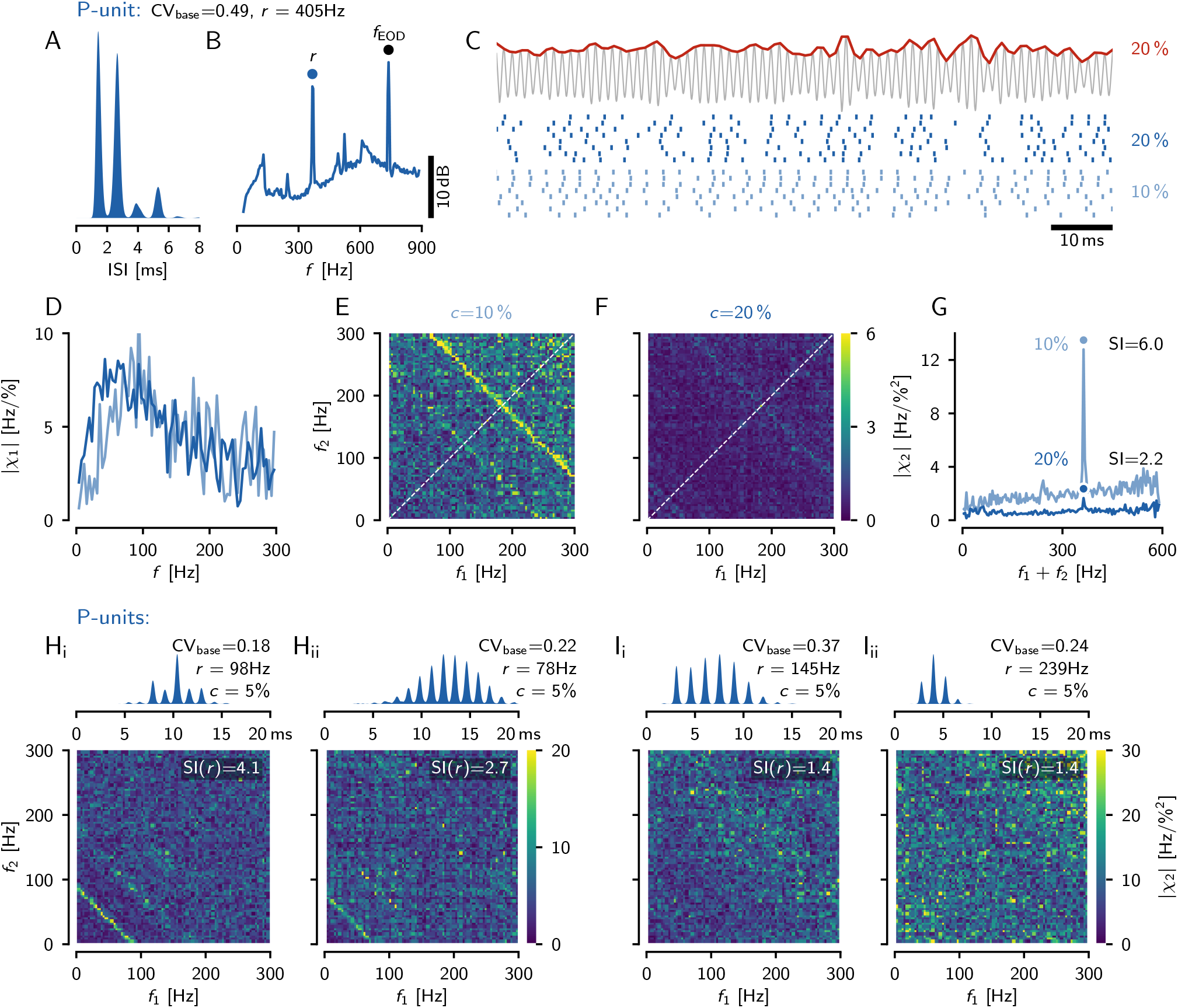
Linear and nonlinear stimulus encoding in example P-units. **A** Interspike interval (ISI) distribution of a cell’s baseline activity, i.e. the cell is driven only by the unperturbed own electric field (cell identifier “2020-10-27-ag”). This cell has a rather high baseline firing rate *r* = 405 Hz and an intermediate CV_base_ = 0.49 of its interspike intervals. **B** Power spectral density of the cell’s baseline response with marked peaks at the cell’s baseline firing rate *r* and the fish’s EOD frequency *f*_EOD_. **C** Random amplitude modulation (RAM) stimulus (top, red, with cutoff frequency of 300 Hz) and evoked responses (spike raster, bottom) of the same P-unit for two different stimulus contrasts (right). The stimulus contrast quantifies the standard deviation of the RAM relative to the fish’s EOD amplitude. **D** Gain of the transfer function (first-order susceptibility), Eq. (4), computed from the responses to 10 % (light blue) and 20 % contrast (dark blue) RAM stimulation of 5 s duration. **E** Absolute value of the second-order susceptibility, Eq. (6), for the low stimulus contrast. An anti-diagonal where the sum of the two stimulus frequencies equals the neuron’s baseline frequency clearly sticks out of the noise floor. **F** At the higher contrast, the anti-diagonal in the absolute value of the second-order susceptibility is much weaker. **G** Second-order susceptibilities projected onto the diagonal (averages over all anti-diagonals of the matrices shown in E, F). The anti-diagonals from E and F show up as a peak close to the cell’s baseline firing rate *r*. The susceptibility index, SI(*r*) Eq. (9), quantifies the height of this peak relative to the values in the vicinity. **H** ISI distributions (top) and second-order susceptibilities (bottom) of two more example P-units (“2021-06-18-ae”, “2017-07-18-ai”) showing an anti-diagonal, but not the full expected triangular structure. **I** Most P-units, however, have a flat second-order susceptibility and consequently their SI(*r*) values are close to one (cell identifiers “2018-08-24-ak”, “2018-08-14-ac”).

Noise stimuli, here random amplitude modulations (RAM) of the EOD (fig. 5 C, top trace, red line), have been commonly used to characterize stimulus-driven responses of sensory neurons using transfer functions (first-order susceptibility), spike-triggered averages, or stimulus-response coherences. Here, we additionally estimate from existing recordings the second-order susceptibility to quantify nonlinear encoding. P-unit spikes align more or less clearly with fluctuations in the RAM stimulus. A higher stimulus intensity, here a higher contrast of the RAM relative to the EOD amplitude (see methods), entrains the P-unit response more clearly (light and dark blue for low and high contrast stimuli, respectively, fig. 5 C). Linear encoding, quantified by the first-order susceptibility or transfer function, Eq. (4), is similar for the two RAM contrasts in this low-CV P-unit (fig. 5 D), as expected for a linear system. The first-order susceptibility is low for low frequencies, peaks in the range below 100 Hz and then falls off again (Benda et al., 2005).

The second-order susceptibility, Eq. (6), quantifies for each combination of two stimulus frequencies *f*_1_ and *f*_2_ the amplitude and phase of the stimulus-evoked response at the sum *f*_1_+*f*_2_ (and also the difference, fig. 1 A). Large values of the second-order susceptibility indicate stimulus-evoked peaks in the response spectrum at the summed frequency that cannot be explained by linear response theory. Similar to the first-order susceptibility, the second-order susceptibility can be estimated directly from the response evoked by a RAM stimulus that stimulates the neuron with a whole range of frequencies simultaneously (figs. 5 E, F). For LIF and theta neuron models driven in the supra-threshold regime, theory predicts nonlinear interactions between the two stimulus frequencies, when the two frequencies *f*_1_ and *f*_2_ or their sum *f*_1_ + *f*_2_ exactly match the neuron’s baseline firing rate *r* (Voronenko and Lindner, 2017; Franzen et al., 2023). Only then, additional stimulus-evoked peaks appear in the spectrum of the spiking response that would show up in the second-order susceptibility as a horizontal, a vertical, and an anti-diagonal line (fig. 1 B).

For the example P-unit, we observe a ridge of elevated second-order susceptibility for the low RAM contrast at *f*_1_ + *f*_2_ = *r* (yellowish anti-diagonal, fig. 5 E). This structure is less prominent for the stronger stimulus (fig. 5 F). Further, the overall level of the second-order susceptibility is reduced with increasing stimulus strength. To quantify the structural changes in the susceptibility matrices we projected the susceptibility values onto the diagonal (white dashed line) by averaging over the anti-diagonals (fig. 5 G). At low RAM contrast this projection indeed has a distinct peak close to the neuron’s baseline firing rate (fig. 5 G, dot on top line). For the higher RAM contrast this peak is much smaller and the overall level of the second-order susceptibility is reduced (fig. 5 G). The reason behind this reduction is that a RAM with a higher contrast is not only a stimulus with an increased amplitude, but also increases the total noise in the system. Increased noise is known to linearize signal transmission (Longtin, 1993; Chialvo et al., 1997; Roddey et al., 2000; Voronenko and Lindner, 2017) and thus the second-order susceptibility is expected to decrease.

Overall we observed in 17 % of the 159 P-units ridges in the second-order susceptibility where the stimulus frequencies add up to the unit’s baseline firing rate. Two more examples are shown in fig. 5 H. However, we never observed the full triangular structure expected from theory (fig. 1 B). In all other P-units, we did not observe any structure in the second-order susceptibility (fig. 5 I).

### Ampullary afferents exhibit strong nonlinear interactions

Electric fish possess an additional electrosensory system, the passive or ampullary electrosensory system, that responds to low-frequency exogenous electric stimuli. The population of ampullary afferents is much less heterogeneous, and known for the much lower CVs of their baseline ISIs (0.06 *<* CV_base_ *<* 0.22, Grewe et al., 2017). Ampullary cells do not phase-lock to the high-frequency EOD and the ISIs have an unimodal distribution (fig. 6 A). As a consequence of the high regularity of their baseline spiking activity, the corresponding power spectrum shows distinct peaks at the baseline firing rate *r* and its harmonics. Since the cells do not respond to the self-generated EOD, there is no sharp peak at *f*_EOD_ (fig. 6 B). When driven by a band-limited white noise stimulus (note: for ampullary afferents this is not an AM stimulus, fig. 6 C), ampullary afferents exhibit very pronounced ridges in the second-order susceptibility, where *f*_1_ + *f*_2_ is equal to *r* or its harmonics (yellow anti-diagonals in fig. 6 E–H), implying strong nonlinear response components at these frequency combinations (fig. 6 G, top). With higher stimulus contrasts these bands get weaker (fig. 6 B), the projection onto the diagonal loses its distinct peak at *r*, and its overall level is reduced (fig. 6 G, bottom). Some ampullary afferents (27 % of 30 afferents), however, do not show any such structure in their second-order susceptibility (fig. 6 I).

**Figure 6.**
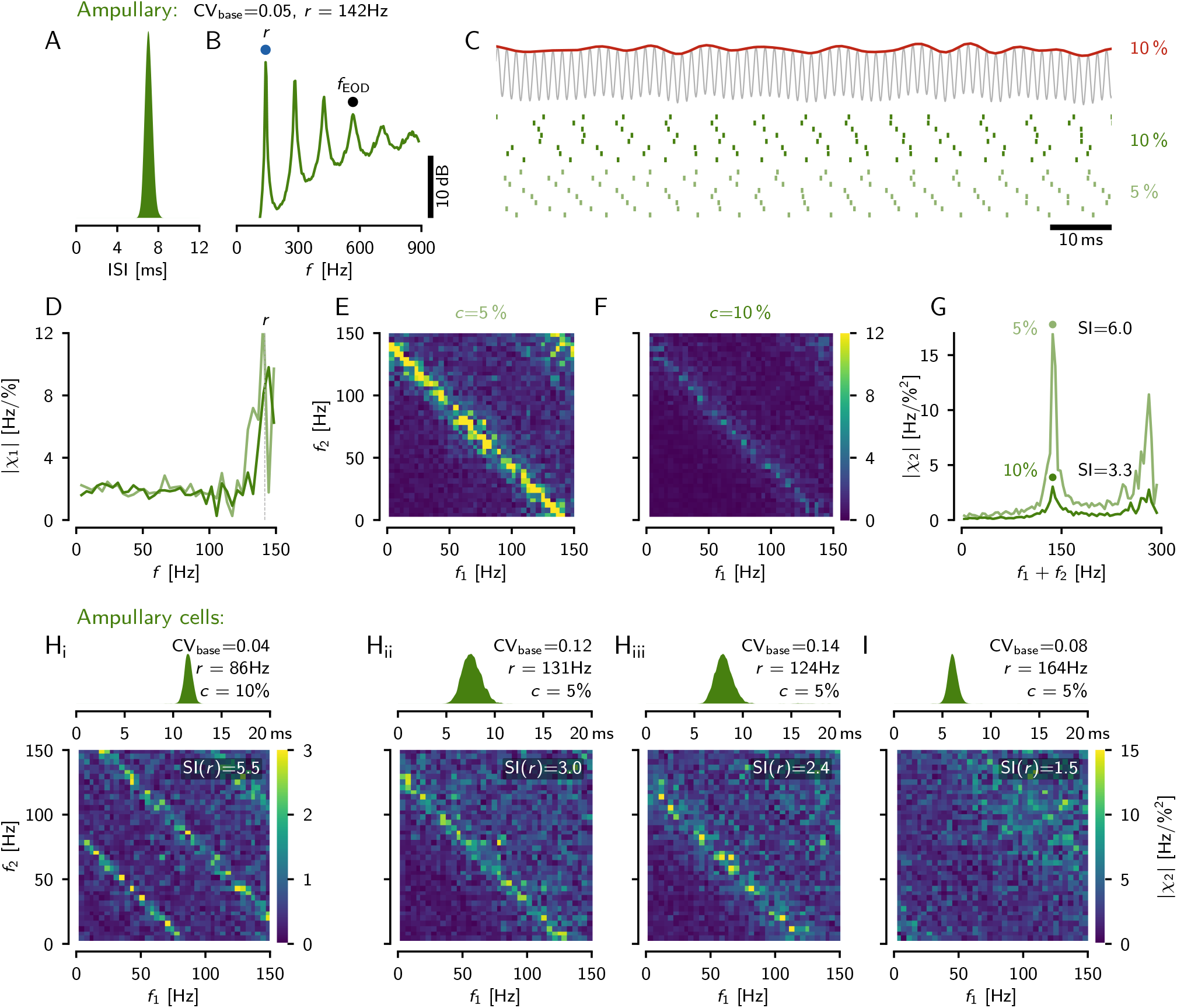
Linear and nonlinear stimulus encoding in example ampullary afferents. **A** Interspike interval (ISI) distribution of the cell’s baseline activity (cell identifier “2012-05-15-ac”). The very low CV of the ISIs indicates almost perfect periodic spiking. **B** Power spectral density of baseline activity with peaks at the cell’s baseline firing rate and its harmonics. Ampullary afferents do not respond to the fish’s EOD frequency, *f*_EOD_ — a sharp peak at *f*_EOD_ is missing. **C** Band-limited white noise stimulus (top, red, with a cutoff frequency of 150 Hz) added to the fish’s self-generated electric field (no amplitude modulation!) and spike raster of the evoked responses (bottom) for two stimulus contrasts as indicated (right). **D** Gain of the transfer function, Eq. (4), of the responses to stimulation with 5 % (light green) and 10 % contrast (dark green) of 10 s duration. **E, F** Absolute value of the second-order susceptibility, Eq. (6), for both stimulus contrasts as indicated. Both show a clear anti-diagonal where the two stimulus frequencies add up to the afferent’s baseline firing rate. **G** Projections of the second-order susceptibilities in E, F onto the diagonal. **H** ISI distributions (top) and second-order susceptibilities (bottom) of three more example afferents with clear anti-diagonals (“2010-11-26-an”, “2010-11-08-aa”, “2011-02-18-ab”). **I** Some ampullary afferents do not show any structure in their second-order susceptibility (“2014-01-16-aj”).

### Model-based estimation of the second-order susceptibility

In the example recordings shown above (figs. 5 and 6), we only observe nonlinear responses where the sum of the two stimulus frequencies matches the neuron’s baseline firing rate, but not where either of the frequencies alone matches the baseline rate. In the following, we investigate this discrepancy to the theoretical expectations (fig. 1, Voronenko and Lindner, 2017; Franzen et al., 2023).

One simple reason could be the lack of data, i.e. the estimation of the second-order susceptibility is not good enough. Electrophysiological recordings are limited in time, and therefore only a limited number of trials, here repeated presentations of the same frozen RAM stimulus, are available. In our data set we have 1 to 199 trials (median: 10) of RAM stimuli with a duration ranging from 2 to 50 s (median: 10 s), total stimulation durations per cell range between 30 and 400 s. Using a temporal resolution of 0.5 ms and FFT segments of 512 samples this yields 105 to 1520 available FFT segments for a specific RAM stimulus. As a consequence, the cross-spectra, Eq. (5), are insufficiently averaged and the full structure of the second-order susceptibility might be hidden in finite-data noise. This experimental limitation can be overcome by using a computational model for the P-unit, a stochastic leaky integrate-and-fire model with adaptation current, dendritic preprocessing, and parameters fitted to the experimentally recorded P-unit (fig. 2) (Barayeu et al., 2023). The model faithfully reproduces the second-order susceptibility of the P-unit estimated from the same low number of FFT (fast fourier transform) segments as in the experiment (*N* = 100, compare faint anti-diagonal in the bottom left corner of the second-order susceptibility in A_ii_ and B_ii_ in fig. 7).

**Figure 7.**
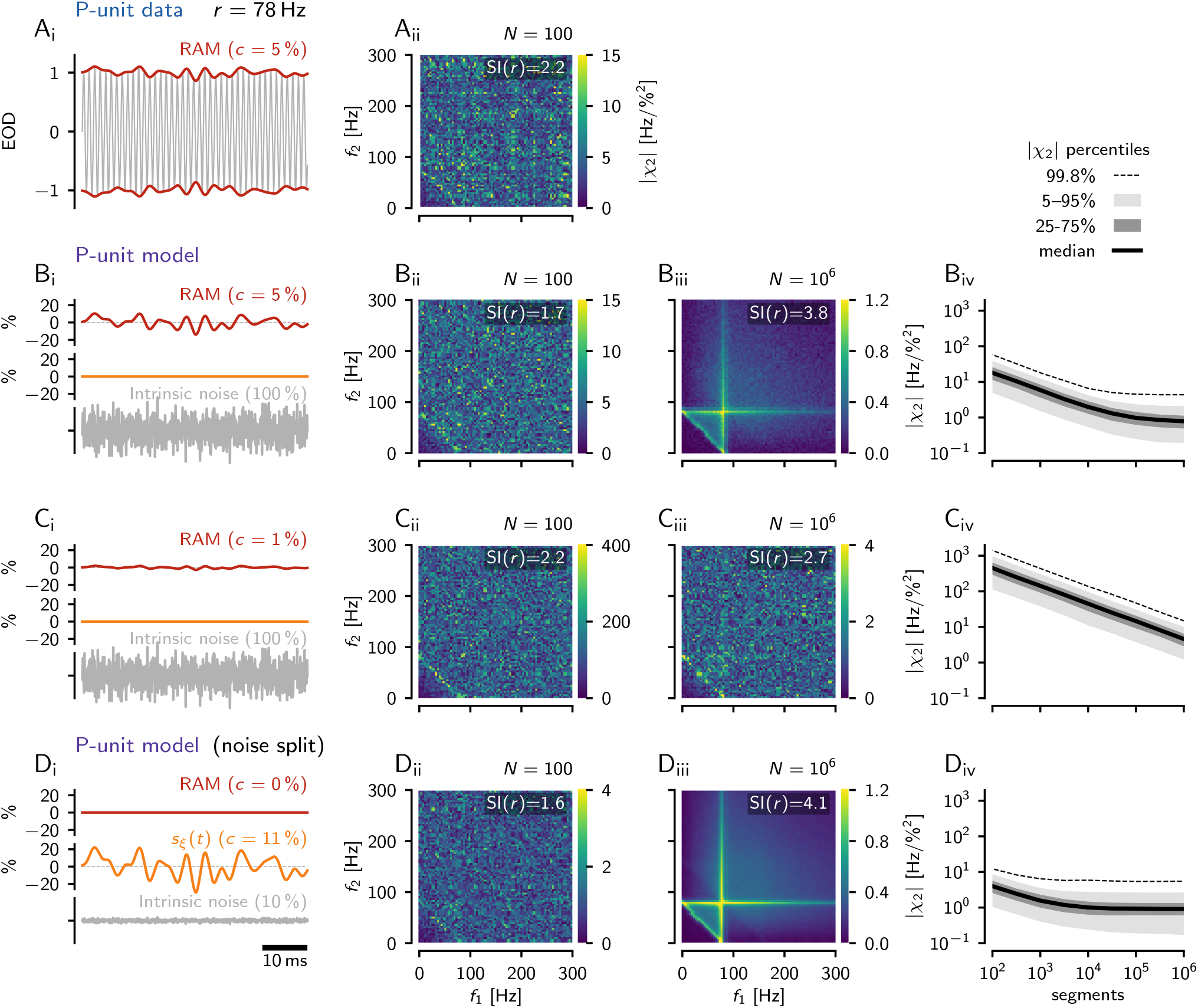
Estimation of second-order susceptibilities. **A** |*χ*_2_(*f*_1_, *f*_2_) | (right) estimated from *N* = 100 256 ms long FFT segments of an electrophysiological recording of another P-unit (cell “2017-07-18-ai”, *r* = 78 Hz, CV_base_ = 0.22) driven with a RAM stimulus with contrast 5 % (left). **B**_i_ *Standard condition* of model simulations with intrinsic noise (bottom) and a RAM stimulus (top). **B**_ii_ |*χ*_2_(*f*_1_, *f*_2_)| estimated from simulations of the cell’s LIF model counterpart (cell “2017-07-18-ai”) based on the same RAM contrast and number of *N* = 100 FFT segments. As in the electrophysiological recording only a weak anti-diagonal is visible. **B**_iii_ Same as B_ii_ but using 10^6^ FFT segments. Now, the expected triangular structure is revealed. **B**_iv_ Convergence of the |*χ*_2_(*f*_1_, *f*_2_) | estimate as a function of FFT segments. **C** At a lower stimulus contrast of 1 % the estimate did not converge yet even for 10^6^ FFT segments. The triangular structure is not revealed yet. **D**_i_ Same as in B_i_ but in the *noise split* condition: there is no external RAM signal (red) driving the model. Instead, a large part (90 %) of the total intrinsic noise is treated as a signal and is presented as an equivalent amplitude modulation (*s*_*ξ*_(*t*), orange, 10.6 % contrast), while the intrinsic noise is reduced to 10 % of its original strength (bottom, see methods for details). **D**_i_ 100 FFT segments are still not sufficient for estimating |*χ*_2_(*f*_1_, *f*_2_) |. **D**_iii_ Simulating one million segments reveals the full expected triangular structure of the second-order susceptibility. **D**_iv_ In the noise-split condition, the |*χ*_2_(*f*_1_, *f*_2_) | estimate converges already at about 10^4^ FFT segments.

In model simulations we can increase the number of FFT segments beyond what would be experimentally possible, here to one million (fig. 7 B_iii_). Then, the estimate of the second-order susceptibility indeed improves. It gets less noisy, the diagonal at *f*_1_ + *f*_2_ = *r* is emphasized, and the vertical and horizontal ridges at *f*_1_ = *r* and *f*_2_ = *r* are revealed. Increasing the number of FFT segments also reduces the order of magnitude of the susceptibility estimate until close to one million the estimate levels out at a low values (fig. 7 B_iv_).

At a lower stimulus contrast of 1 % (fig. 7 C), however, one million FFT segments are still not sufficient for the estimate to converge (fig. 7 C_iv_). Still only a faint anti-diagonal is visible (fig. 7 C_iii_).

Using a broadband stimulus increases the effective input-noise level. This may linearize signal transmission and suppress potential nonlinear responses (Longtin, 1993; Chialvo et al., 1997; Roddey et al., 2000; Voronenko and Lindner, 2017). Assuming that the intrinsic noise level in this P-unit is small enough, the full expected structure of the second-order susceptibility should appear in the limit of weak AMs. As we just have seen, this cannot be done experimentally. In the model, however, we know the time course of the intrinsic noise and can use this knowledge to determine the susceptibilities by input-output correlations via the Furutsu-Novikov theorem (Furutsu, 1963; Novikov, 1965). This theorem, in its simplest form, states that the cross-spectrum *S*_*xη*_(*ω*) of a Gaussian noise *η*(*t*) driving a nonlinear system and the system’s output *x*(*t*) is proportional to the linear susceptibility according to *S*_*xη*_(*ω*) = *χ*(*ω*)*S*_*ηη*_(*ω*). Here *χ*(*ω*) characterizes the linear response to an infinitely weak signal *s*(*t*) in the presence of the background noise *η*(*t*). Likewise, the nonlinear susceptibility can be determined in an analogous fashion from higher-order input-output cross-spectra (see methods, equations (5) and (6)) (Egerland, 2021). In line with an alternative derivation of the Furutsu-Novikov theorem (Lindner, 2022), we can split the total noise and consider a fraction of it as a stimulus. This allows us to calculate the susceptibility from the cross-spectrum between the output and this stimulus fraction of the noise. Adapting this approach to our P-unit model (see methods), we replace the intrinsic noise by an approximately equivalent RAM stimulus *s*_*ξ*_(*t*) and a weak remaining intrinsic noise 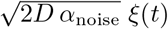 with *α*_noise_ = 0.1 (see methods, equations (18), (20), (19), fig. 7 D_i_). We tuned the amplitude of the RAM stimulus *s*_*ξ*_(*t*) such that the output firing rate and variability (CV of interspike intervals) are the same as in the baseline activity (i.e. full intrinsic noise 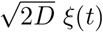 in the voltage equation but no RAM) and compute the cross-spectra between the RAM part of the noise *s*_*ξ*_(*t*) and the output spike train. This procedure has two consequences: (i) by means of the cross-spectrum between the output and *s*_*ξ*_(*t*), which is a large fraction of the noise, the signal-to-noise ratio of the measured susceptibilities is drastically improved and thus the estimate converges already at about ten thousand FFT segments (fig. 7 D_iv_); (ii) the total noise in the system has been reduced (by what was before the external RAM stimulus *s*(*t*)), which makes the system more nonlinear. For both reasons we now see the expected nonlinear features in the second-order susceptibility for a sufficient number of FFT segments (fig. 7 D_iii_), but not for a number of segments comparable to the experiment (fig. 7 D_ii_). In addition to the strong response at *f*_1_ + *f*_2_ = *r*, we now also observe pronounced nonlinear responses at *f*_1_ = *r* and *f*_2_ = *r* (vertical and horizontal lines, fig. 7 D_iii_).

### Weakly nonlinear interactions in many model cells

In the previous section we have shown one example cell for which we find in the corresponding model the expected strong ridges in the second-order susceptibility (fig. 7 B_iii_,D_iii_). Using our 39 P-unit models, we now can explore how many P-unit model neurons show such a triangular structure.

By just looking at the second-order susceptibilities estimated using the noise-split method (first column of fig. 8) we can readily identify strong triangular patterns in 11 of the 39 model cells (28 %, see fig. 8 A_i_ & B_i_ for two examples). In another 5 cells (13 %) the triangle is much weaker and sits on top of a smooth bump of elevated second-order susceptibility (fig. 8 C_i_ shows an example). The remaining 23 model cells (59 %) show no triangle (see fig. 8 D_i_ for an example).

**Figure 8.**
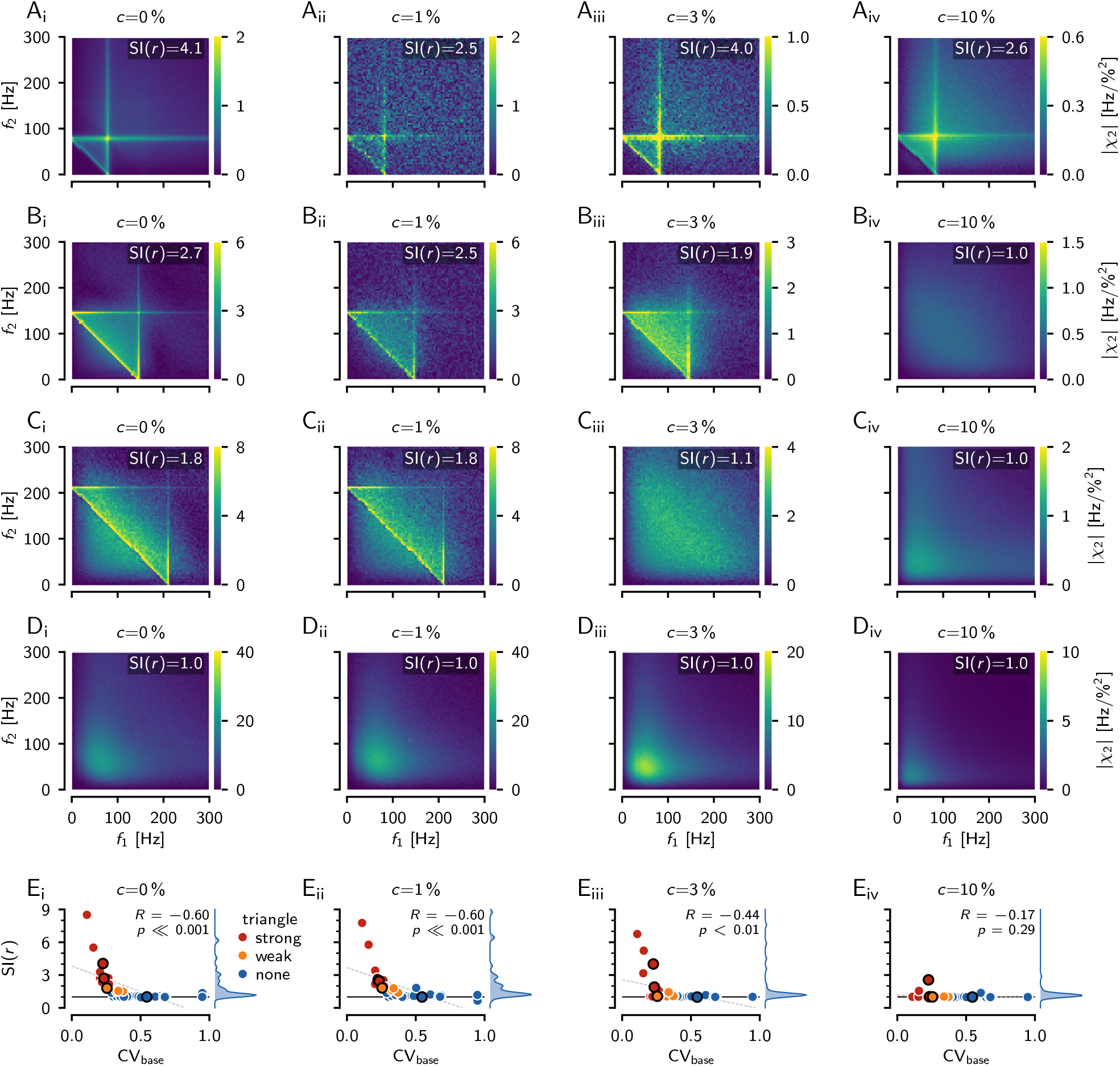
Dependence of second order susceptibility on stimulus contrast. **A** Second-order susceptibilities estimated for increasing stimulus contrasts of *c* = 0, 1, 3 and 10 % as indicated (*N* = 10^7^ FFT segments for *c* = 1 %, *N* = 10^6^ segments for all other contrasts). *c* = 0 % refers to the noise-split configuration (limit to vanishing external RAM signal, see fig. 7 D). Shown are simulations of the P-unit model cell “2017-07-18-ai”) with a baseline firing rate of 82 Hz and CV_base_ = 0.23. The cell shows a clear triangular pattern in its second-order susceptibility even up to a contrast of 10 %. Note, that for *c* = 1 % (D_ii_), the estimate did not converge yet. **B** Cell “2012-12-13-ao” (baseline firing rate of 146 Hz, CV= 0.23) also has strong interactions at its baseline firing rate that survive up to a stimulus contrast of 3 %. **C** Model cell “2012-12-20-ac” (baseline firing rate of 212 Hz, CV= 0.26) shows a weak triangular structure in the second-order susceptibility that vanishes for stimulus contrasts larger than 1 %. **D** Cell “2013-01-08-ab” (baseline firing rate of 218 Hz, CV= 0.55) does not show any triangular pattern in its second-order susceptibility. Nevertheless, interactions between low stimulus frequencies become substantial at higher contrasts. **E** The presence of an elevated second-order susceptibility where the stimulus frequency add up to the neuron’s baseline frequency, can be identified by the susceptibility index (SI(*r*), Eq. (9)) greater than one (horizontal black line). The SI(*r*) (density to the right) is plotted as a function of the model neuron’s baseline CV for all 39 model cells. Model cells have been visually categorized based on the strong (11 cells) or weak (5 cells) presence of a triangular pattern in their second-order susceptibility estimated in the noise-split configuration (legend). The cells from A–D are marked by black circles. Pearson’s correlation coefficients *r*, the corresponding significance level *p* and regression line (dashed gray line) are indicated. The higher the stimulus contrast, the fewer cells show weakly nonlinear interactions as expressed by the triangular structure in the second-order susceptibility.

This categorization is supported by the susceptibility index, SI(*r*), Eq. (9), which quantifies the height of the ridge where the stimulus frequencies add up to the neuron’s baseline firing rate relative to the background. Values above one indicate an elevated ridge. The absence of such a ridge results in values close to one. Indeed, the cells showing only a weak triangle (orange) arise out of values around one and the cells showing strong triangles (red) have consistently SI(*r*) values exceeding 1.8 (fig. 8 E_i_).

The SI(*r*) correlates with the CVs of the cell’s baseline interspike intervals (*r* = − 0.60, *p <* 0.001). The lower the cell’s CV_base_, the higher the SI(*r*) value and thus the stronger the triangular structure of its second-order susceptibility. The model cells with the most distinct triangular pattern in their second-order susceptibility are the ones with the lowest CVs, hinting at low intrinsic noise levels.

### Weakly nonlinear interactions vanish for higher stimulus contrasts

As pointed out above, noise stimuli act as a linearizing background noise and thus may shift the neural signal transmission into a regime free of nonlinearities (see discussion). Sine-wave stimuli, as for example used in the introductory figure 4, do not have such a linearizing effect. At which amplitudes a noise stimulus effectively linearizes the system, also depends on cellular properties of the neuron, or on the parameters in our leaky integrate-and-fire neuron. This is what we explore now by varying the contrast of RAM stimuli driving our model cells.

In the model cells we estimated second-order susceptibilities for RAM stimuli with a contrast of 1, 3, and 10 %. The estimates for 1 % contrast (fig. 8 E_ii_) were quite similar to the estimates from the noise-split method, corresponding to a stimulus contrast of 0 % (*r* = 0.97, *p* « 0.001). Thus, RAM stimuli with 1 % contrast are sufficiently small to not destroy weakly nonlinear interactions by their linearizing effect. At this low contrast, 51 % of the model cells have an SI(*r*) value greater than 1.2.

At a RAM contrast of 3 % the SI(*r*) values become smaller (fig. 8 E_iii_). Only 7 cells (18 %) have SI(*r*) values exceeding 1.2. Finally, at 10 % the SI(*r*) values of all cells drop below 1.2, except for three cells (8 %, fig. 8 E_iv_). The cell shown in fig. 8 B is one of them. At 10 % contrast the SI(*r*) values are no longer correlated with the ones in the noise-split configuration (*r* = 0.32, *p* = 0.05). To summarize, the regime of distinct nonlinear interactions at frequencies matching the baseline firing rate extends in this set of P-unit model cells to stimulus contrasts ranging from a few percents to about 10 %.

### Weakly nonlinear interactions can be deduced from limited data

Estimating second-order susceptibilities reliably requires large numbers (millions) of FFT segments (fig. 7). Electrophysiological measurements, however, suffer from limited recording durations and estimating weakly nonlinear interactions from just a few hundred segments appears futile. To what extend are such limited-data estimates still informative?

The second-order susceptibility matrices that are based on only 100 segments look flat and noisy, lacking the triangular structure (fig. 9 B). The anti-diagonal ridge, however, seems to be present whenever the converged estimate shows a clear triangular structure (compare fig. 9 B and fig. 9 A). Comparing SI(*r*) values based on 100 FFT segments to the ones based on one or ten million segments for all 39 model cells (fig. 9 C) supports this impression. They correlate quite well at contrasts of 1 % and 3 % (*r* = 0.9, *p* « 0.001). At a contrast of 10 % this correlation is weaker (*r* = 0.38, *p* < 0.05), because there are only three cells left with SI(*r*) values greater than 1.2. Despite the good correlations, care has to be taken to set a threshold on the SI(*r*) values for deciding whether a triangular structure would emerge for a much higher number of segments. Because at low number of segments the estimates are noisier, there could be false positives for a too low threshold. Setting the threshold to 1.8 avoids false positives for the price of a few false negatives.

**Figure 9.**
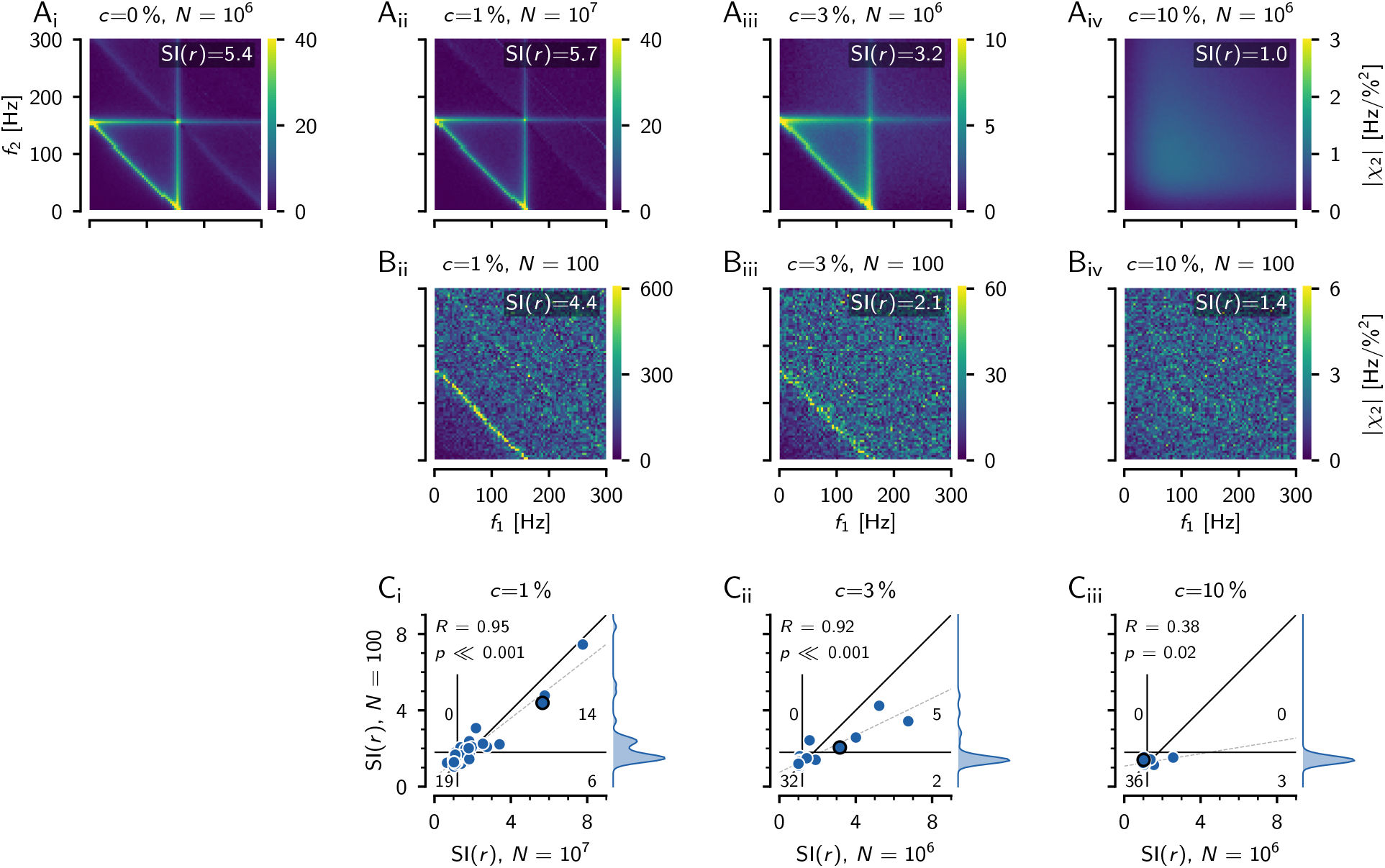
Inferring the triangular structure of the second-order susceptibility from limited data. **A** Reliably estimating the structure of the second-order susceptibility requires a high number of FFT segments *N* in the order of one or even ten millions. As an example, susceptibilities of the model cell “2012-12-21-ak” (baseline firing rate of 157 Hz, CV=0.15) are shown for the noise-split configuration (*c* = 0 %) and RAM stimulus contrasts of *c* = 1, 3, and 10 % as indicated. For contrasts below 10 % this cell shows a nice triangular pattern in its susceptibilities, quite similar to the introductory example of a LIF in fig. 1. **B** However, with limited data of *N* = 100 trials the susceptibility estimates are noisy and show much less structure, except for the anti-diagonal at the cell’s baseline firing rate. The SI(*r*) quantifies the height of this ridge where the two stimulus frequencies add up to the neuron’s baseline firing rate. **C** Correlations between the estimates of SI(*r*) based on 100 FFT segments (density to the right) with the converged ones based on one or ten million segments at a given stimulus contrast for all *n* = 39 model cells. The black circle marks the model cell shown in A and B. The black diagonal line is the identity line and the dashed line is a linear regression. The correlation coefficient and corresponding significance level are indicated in the top left corner. The thin vertical line is a threshold at 1.2, the thin horizontal line a threshold at 1.8. The number of cells within each of the resulting four quadrants denote the false positives (top left), true positives (top right), true negatives (bottom left), and false negatives (bottom right) for predicting a triangular structure in the converged susceptibility estimate from the estimates based on only 100 segments.

Overall, observing SI(*r*) values greater than about 1.8, even for a number of FFT segments as low as one hundred, seems to be a reliable indication for a triangular structure in the second-order susceptibility at the corresponding stimulus contrast. Small stimulus contrasts of 1 % are less informative, because of their bad signal-to-noise ratio. Intermediate stimulus contrasts around 3 % seem to be optimal, because there, most cells still have a triangular structure in their susceptibility and the signal-to-noise ratio is better. At RAM stimulus contrasts of 10 % or higher the signal-to-noise ratio is even better, but only few cells remain with weak triangularly shaped susceptibilities that might be missed as a false positives.

### Low CVs and weak responses predict weakly nonlinear responses

Now we are prepared to evaluate our pool of 39 P-unit model cells, 172 P-units, and 30 ampullary afferents recorded in 80 specimen of *Apteronotus leptorhynchus*. For direct comparison across cells we condensed the structure of the second-order susceptibilities into SI(*r*) values, Eq. (9). Both, model and experimental SI(*r*) estimates, Eq. (9), are based on 100 FFT segments.

In the P-unit models, each model cell contributed with three RAM stimulus presentations with contrasts of 1, 3, and 10 %, resulting in *n* = 117 data points. 19 (16 %) had SI(*r*) values larger than 1.8, indicating the expected ridges at the baseline firing rate in their second-order susceptibility. The lower the cell’s baseline CV, i.e. the less intrinsic noise, the higher the SI(*r*) (fig. 10 A_i_).

**Figure 10.**
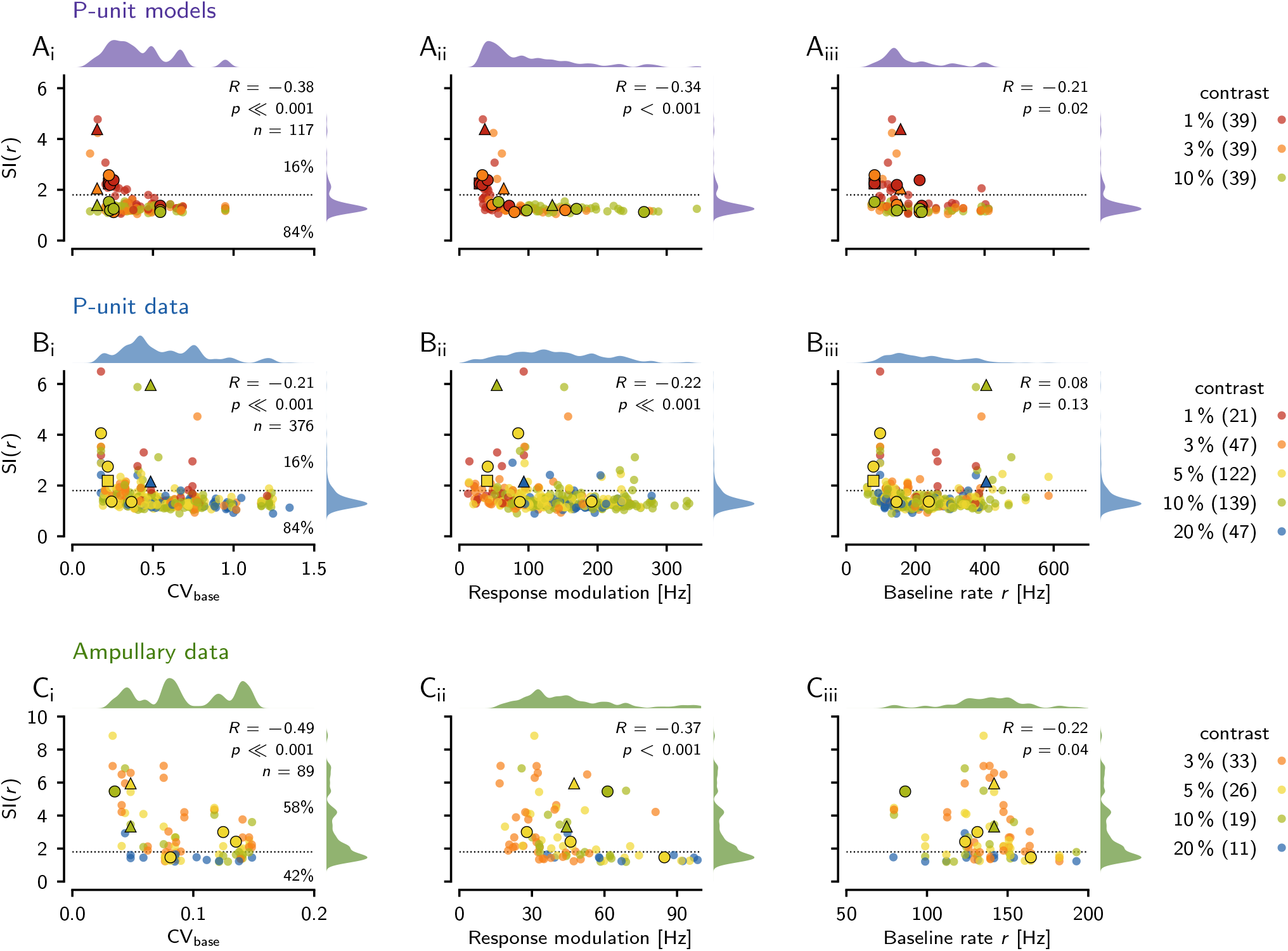
Nonlinear responses in P-units and ampullary afferents. The second-order susceptibility is condensed into the susceptibility index, SI(*r*) Eq. (9), that quantifies the relative amplitude of the ridge where the two stimulus frequencies add up to the cell’s baseline firing rate *r* (see fig. 5 G). In both the models and the experimental data, the SI(*r*) was estimated based on 100 FFT segments. The SI(*r*) is plotted against the cells’ CV of its baseline interspike intervals (left column), the response modulation (standard deviation of firing rate evoked by the band-limited white-noise stimulus) — a measure of effective stimulus strength (center column), and the cell’s baseline firing rate (right column). Pearson’s correlation coefficient *R*, the corresponding significance level *p* and the number of samples *n* are indicated. Kernel-density estimates of the distributions of the displayed quantities are plotted on top and right. Data points are color coded by stimulus contrasts, which are listed together with the corresponding number of samples in the legend to the right. The horizontal dashed line marks a threshold for SI(*r*) values at 1.8 and the percentages to the right denote the fractions of samples above and below this threshold. **A** The SI(*r*) of all 39 model P-units measured with RAM stimuli with a cutoff frequency of 300 Hz. The black square marks the cell from fig. 7 C, the circles the four cells shown in fig. 8 A–D, and the triangle the cell from fig. 9 A–B. **B** Electrophysiological data from 172 P-units. Each cell contributes on average with 2 (min. 1, max. 10) RAM stimulus presentations to the *n* = 376 data points. The RAMs had cutoff frequencies of 300 Hz (352 samples) and 400 Hz (24 samples). The two black triangles mark the responses of the example P-unit from fig. 5 E,F, the circles the other four examples from fig. 5 H, and the triangle the unit from fig. 7 A. **C** Recordings from 30 ampullary afferents, each contributing on average 3 (min. 1, max. 7) RAM stimulus presentations to *n* = 89 data points. Stimuli had a cutoff frequency of 150 Hz. The two black triangles mark the responses of the example ampullary afferent from fig. 6 E,F, and the circles the other four examples from fig. 6 H.

The effective stimulus strength also plays a role in predicting the SI(*r*) values. We quantify the effect of stimulus strength on a cell’s response by the response modulation — the standard deviation of a cell’s firing rate in response to a RAM stimulus. The lower the response modulation, i.e. the weaker the effective stimulus, the higher the S(*r*) (fig. 10 A_ii_). Although there is a tendency of low stimulus contrasts to evoke lower response modulations, response modulations evoked by each of the three contrasts overlap substantially, emphasizing the strong heterogeneity of the P-units’ sensitivity (Grewe et al., 2017). Cells with high SI(*r*) values are the ones with baseline firing rate below 200 Hz (fig. 10 A_iii_).

In comparison to the experimentally measured P-unit recordings, the model cells are skewed to lower baseline CVs (Mann-Whitney *U* = 13986, *p* = 3 × 10^*−*9^), because the models are not able to reproduce bursting, which we observe in many P-units and which leads to high CVs. Also the response modulation of the models is skewed to lower values (Mann-Whitney *U* = 14051, *p* = 4 × 10^*−*9^), because in the measured cells, response modulation is positively correlated with baseline CV (Pearson *R* = 0.34, *p* = 1 × 10^*−*4^), i.e. bursting cells are more sensitive. Median baseline firing rate in the models is by 53 Hz smaller than in the experimental data (Mann-Whitney *U* = 17034, *p* = 0.0002).

In the experimentally measured P-units, each of the 172 cells contributes on average with two RAM stimulus presentations, presented at contrasts ranging from 1 to 20 % to the 376 samples. Despite the mentioned differences between the P-unit models and the measured data, the SI(*r*) values do not differ between models and data (median of 1.3, Mann-Whitney *U* = 19702, *p* = 0.09) and also 16 % of the samples from all presented stimulus contrasts exceed the threshold of 1.8. The SI(*r*) values of the P-unit population correlate weakly with the CV of the baseline ISIs that range from 0.18 to 1.35 (median 0.49). Cells with lower baseline CVs tend to have more pronounced ridges in their second-order susceptibilities than those with higher baseline CVs (fig. 10 B_i_).

Samples with weak responses to a stimulus, due to low sensitivity or a weak stimulus, have higher SI(*r*) values in comparison to strongly responding cells, most of them having flat second-order susceptibilities (fig. 10 B_ii_). P-units with low or high baseline firing rates can have large SI(*r*) (fig. 10 B_iii_). How pronounced nonlinear response components are in P-units thus depends on the baseline CV (a proxy for the internal noise level), and the response strength during stimulation (effective output noise).

The population of ampullary cells is more homogeneous, with generally lower baseline CVs than P-units (Mann-Whitney *U* = 33464, *p* = 9 × 10^*−*49^). Accordingly, SI(*r*) values of ampullary cells (median 2.3) are indeed higher than in P-units (median 1.3, Mann-Whitney *U* = 6450, *p* = 2 × 10^*−*19^). 52 samples (58 %) with SI(*r*) values greater than 1.8 would have a triangular structure in their second-order susceptibilities.

Ampullary cells also show a negative correlation with baseline CV, despite their narrow distribution of CVs ranging from 0.03 to 0.15 (median 0.09) (fig. 10 C_i_). Again, sensitive cells with stronger response modulations are at the bottom of the SI(*r*) distribution with values close to one (fig. 10 C_ii_). Similar to P-units, the baseline firing rate does not predict SI(*r*) values (fig. 10 C_iii_).

## Discussion

Theoretical work (Voronenko and Lindner, 2017; Franzen et al., 2023) studied analytically and numerically the weakly-nonlinear responses of spike generating LIF and theta model neurons driven by two sine waves with distinct frequencies. We here investigated such nonlinear responses in two types of electroreceptor afferents that differ in their intrinsic noise levels (Grewe et al., 2017) using band-limited white-noise stimuli to estimate second-order susceptibilities. Following Voronenko and Lindner (2017) we expected to observe distinct ridges in the second-order susceptibility where either of the stimulus frequencies alone or their sum matches the baseline firing rate. We find traces of these nonlinear responses in the majority of ampullary afferents. In P-units, however, only a minority of the recorded cells, i.e. those characterized by low intrinsic noise levels and low output noise, show signs of such nonlinear responses. Complementary model simulations demonstrate in the limit of high numbers of FFT segments, that the estimates from the electrophysiological data are indeed indicative of the theoretically expected triangular structure of supra-threshold weakly nonlinear responses. With this, we provide evidence for weakly-nonlinear responses of a spike generator at low intrinsic noise levels or low stimulus amplitudes in real sensory neurons.

### Intrinsic noise limits nonlinear responses

The weakly nonlinear regime with its triangular pattern of elevated second-order susceptibility resides between the linear and a stochastic mode-locking regime. Too strong intrinsic noise linearizes the system and wipes out the structure of the second-order susceptibility (Voronenko and Lindner, 2017, fig. 1 B). The CV of the baseline interspike interval is a proxy for the intrinsic noise in the cells (Vilela and Lindner, 2009, note however the effect of coherence resonance for excitable systems close to a bifurcation, Pikovsky and Kurths, 1997; Lindner et al., 2004). In both cell types, we observe a negative correlation between the second-order susceptibility at *f*_1_ + *f*_2_ = *r* and the baseline CV (fig. 10), indicating that it is the level of intrinsic noise that shapes nonlinear responses. Still, only 18 % of the P-units analyzed in this study show relevant nonlinear responses. On the other hand, the majority (74 %) of the ampullary cells show nonlinear responses as they have generally lower CVs (median of 0.09).

These findings are in line with previous theoretical and experimental studies showing the linearizing effects of noise in other nonlinear systems (Roddey et al., 2000; Chialvo et al., 1997). Increased intrinsic noise has been demonstrated to increase the CV and to reduce nonlinear phase-locking in vestibular afferents (Schneider et al., 2011). Reduced noise, on the other hand, has been associated with stronger nonlinearity in pyramidal cells of the ELL (Chacron, 2006). Only in cells with sufficiently low levels of intrinsic noise, weakly nonlinear responses can be observed.

### Linearization by white-noise stimulation

Not only the intrinsic noise but also the stimulation with external white-noise linearizes the cells. This applies to both, the stimulation with AMs in P-units (fig. 10 B_ii_) and direct stimulation in ampullary cells (fig. 10 C_ii_). The stronger the effective stimulus, the less pronounced are the ridges in the second-order susceptibility (see fig. 5 E&F for a P-unit example and fig. 6 E&F for an ampullary cell). This linearizing effect of noise stimuli limits the weakly nonlinear regime to small stimulus amplitudes. At higher stimulus amplitudes, however, other nonlinearities of the system eventually show up in the second-order susceptibility.

In order to characterize weakly nonlinear responses of the cells in the limit to vanishing stimulus amplitudes we utilized the Furutsu-Novikov theorem (Novikov, 1965; Furutsu, 1963). Following Lindner (2022), a substantial part of the intrinsic noise of a P-unit model (Barayeu et al., 2023) is treated as signal. Performing this noise-split trick we can estimate the weakly nonlinear response without the linearizing effect of an additional external white noise stimulus. 41 % of the model cells then show the full nonlinear structure (fig. 8 E_i_) known from analytical derivations and simulations of basic LIF and theta models driven with pairs of sine-wave stimuli (Voronenko and Lindner, 2017; Franzen et al., 2023). Previous studies on second-order nonlinearities have not observed the weakly nonlinear regime, probably because of the linearizing effects of strong noise stimuli (Victor et al., 1977; Schanze and Eckhorn, 1997; Neiman and Russell, 2011a).

### Characterizing nonlinear coding from limited experimental data

Estimating the Volterra series from limited experimental data is usually restricted to the first two or three orders, which might not be sufficient for a proper prediction of the neuronal response (French et al., 2001). As we have demonstrated, a proper estimation of just the second-order susceptibility in the weakly nonlinear regime is challenging in electrophysiological experiments (fig. 9). Our minimum of 100 FFT segments corresponds to just 26 s of stimulation and thus is not a challenge. However, the estimates of the second-order susceptibilities start to converge only beyond 10 000 FFT segments, corresponding to 43 min of stimulation. Often, however, more than one million segments (corresponding to 71 hours of recording), were needed, which is clearly out of reach. We have demonstrated that even in non-converged estimates based on short recordings, the presence of a anti-diagonal ridge is sufficient to predict a triangular pattern in a converged estimate.

Making assumptions about the nonlinearities in a system also reduces the amount of data needed for parameter estimation. In particular, models combining linear filtering with static nonlinearities (Chichilnisky, 2001) have been successful in capturing functionally relevant neuronal computations in visual (Gollisch and Meister, 2009) as well as auditory systems (Clemens and Ronacher, 2013). Linear methods based on backward models for estimating the stimulus from neuronal responses, have been extensively used to quantify information transmission in neural systems (Theunissen et al., 1996; Borst and Theunissen, 1999; Wessel et al., 1996; Machens et al., 2001), because backward models do not need to generate action potentials that involve strong nonlinearities (Rieke et al., 1999).

### Nonlinear encoding in ampullary cells

The afferents of the passive electrosensory system, the ampullary cells, exhibit strong second-order susceptibilities (fig. 10). Ampullary cells more or less directly translate external low-frequency electric fields into afferent spikes, much like in the standard LIF and theta models used by Lindner and colleagues (Voronenko and Lindner, 2017; Franzen et al., 2023). Indeed, we observe in the ampullary cells similar second-order nonlinearities as in LIF models.

Ampullary stimuli originate from the muscle potentials induced by prey movement (Kalmijn, 1974; Engelmann et al., 2010; Neiman and Russell, 2011b). For a single prey item such as *Daphnia*, these potentials are often periodic but the simultaneous activity of a swarm of prey resembles Gaussian white noise (Neiman and Russell, 2011b). Linear and nonlinear encoding in ampullary cells has been studied in great detail in the paddlefish (Neiman and Russell, 2011b). The power spectrum of the baseline response shows two main peaks: One peak at the baseline firing frequency, a second one at the oscillation frequency of primary receptor cells in the epithelium, plus interactions of both. Linear encoding in the paddlefish shows a gap at the epithelial oscillation frequency, instead, nonlinear responses are very pronounced there.

Ampullary stimulus encoding is somewhat different in *A. leptorhynchus*. The power spectrum of the spontaneous response is dominated by only the baseline firing rate and its harmonics, a second oscillator is not visible. The baseline firing frequency, however, is outside the linear coding range (Grewe et al., 2017) while it is within the linear coding range in paddlefish (Neiman and Russell, 2011b). Interestingly, the nonlinear response in the paddlefish ampullary cells increases with stimulus intensity while it disappears in our case (fig. 10 C_ii_) indicating that paddlefish data have been recorded above the weakly-nonlinear regime.

The population of ampullary cells is very homogeneous with respect to the baseline rate (131 ± 29 Hz) and stimulus encoding properties (Grewe et al., 2017). This implies that, if the stimulus contains the appropriate frequency components that sum up to the baseline rate, the resulting nonlinear response appears at the baseline rate that is similar in the full population of ampullary cells and that is outside the linear coding range. Postsynaptic cells integrating ampullary input might be able to extract such nonlinear responses. How such nonlinear effects might influence prey detection should be addressed in future studies.

### Nonlinear encoding in P-units

In contrast to the ampullary cells, P-units respond to the amplitude modulation of the self-generated EOD. Extracting the AM requires a (threshold) nonlinearity (Middleton et al., 2006; Stamper et al., 2012; Savard et al., 2011; Barayeu et al., 2023). This nonlinearity, however, does not show up in our estimates of the susceptibilities, because in our analysis we directly relate the AM waveform to the recorded cellular responses. Encoding the time-course of the AM, however, has been shown to be linear over a wide range of AM amplitudes and frequencies (Xu et al., 1996; Benda et al., 2005; Gussin et al., 2007; Grewe et al., 2017; Savard et al., 2011). In contrast, we here have demonstrated nonlinear interactions originating from the spike generator for broad-band noise stimuli with small amplitudes and for stimulation with two distinct frequencies. Both settings have not been studied yet.

Noise stimuli have the advantage that a range of frequencies can be measured with a single stimulus presentation and they have been successfully applied to characterize sensory coding in many systems (French and Butz, 1973; Marmarelis et al., 1999; Borst and Theunissen, 1999; Chacron et al., 2005; Grewe et al., 2017). The natural stimuli encoded by P-units are, however, periodic amplitude modulations of the self-generated electric field which arise from the superposition of the own and foreign EODs. Such interactions usually occur between low numbers of close-by fish and thus the AMs are a mixture of a few distinct frequencies with substantial amplitudes (Stamper et al., 2010; Fotowat et al., 2013; Henninger et al., 2020). How informative are the second-order susceptibilities observed under noise stimulation for the encoding of distinct frequencies? Broadband noise stimuli introduce additional noise that linearizes the dynamics of the system. In contrast, a pure sine wave stimulation is spectrally focused and drives the system on the background of the intrinsic noise. This explains why we can observe nonlinear interactions between sine wave stimuli with distinct frequencies and substantial power (fig. 3) although these interactions vanish when stimulating with noise stimuli of similar contrast (fig. 8).

The encoding of secondary AMs or social envelopes that arise from relative movement or the interaction of more than two animals (Stamper et al., 2012) requires another nonlinearity in addition to the one needed for extracting the AM. Initially, this nonlinearity was attributed to downstream processing (Middleton et al., 2006, 2007). Later studies showed that already the electroreceptors can encode such information whenever the firing rate saturates at zero or the maximum rate at the EOD frequency (Savard et al., 2011). Based on our work, we predict that P-units with low CVs encode the social envelopes even under weak stimulation, whenever the resulting beat frequencies match or add up to the baseline firing rate. Then difference frequencies show up in the response spectrum that characterize the slow envelope.

The weakly nonlinear interactions in low-CV P-units could facilitate the detectability of faint signals during three-animal interactions as found in courtship contexts in the wild (Henninger et al., 2018). The detection of a faint, distant intruder male could be improved by the presence of a nearby strong female stimulus, because of the nonlinear interaction terms (Schlungbaum and Lindner, 2023). This boosting effect is, however, very specific with respect to the stimulus frequencies and a given P-unit’s baseline frequency. The population of P-units is very heterogeneous in their baseline firing rates and CVs (50–450 Hz and 0.1–1.4, respectively, fig. 10 B, Grewe et al., 2017; Hladnik and Grewe, 2023). The range of baseline firing rates thus covers substantial parts of the beat frequencies that may occur during animal interactions (Henninger et al., 2018, 2020), while the number of P-units showing weakly nonlinear responses is small. Whether and how this information is specifically maintained and read out by pyramidal cells in the electrosensory lateral line lobe (ELL) in the hindbrain onto which P-units converge (Krahe and Maler, 2014; Maler, 2009) is an open question.

Electric fish are able to slowly modulate their EOD frequency, as for example during the so-called jamming-avoidance-response (Fortune et al., 2020). Such behaviors modify the resulting beat frequency by a few Hertz. This could in principle increase the chance that the now slowly changing beat frequency matches at some point the baseline firing rate of a P-unit, where the weakly nonlinear responses then enhance the detectability of another conspecific (Schlungbaum and Lindner, 2023). Furthermore, transient changes in EOD frequency on timescales of tens of milliseconds up to a few seconds are known as chirps and rises, respectively, and are involved in courtship and aggression behaviors (Henninger et al., 2018; Raab et al., 2021). How the encoding of such transient frequency modulations is affected by the nonlinearities described here is another open question, since the presented analysis focuses on stationary signals.

## Conclusions

We have demonstrated pronounced nonlinear responses in primary electrosensory afferents at weak stimulus amplitudes and sufficiently low intrinsic noise levels. The observed nonlinearities match the expectations from previous theoretical studies (Voronenko and Lindner, 2017; Franzen et al., 2023). The resulting nonlinear components introduce spectral components not present in the original stimulus, that may provide an edge in the context of signal detection problems at stimulus amplitudes close to threshold (Schlungbaum and Lindner, 2023). Electrosensory afferents share an evolutionary history with hair cells (Baker, 2019) and share many response properties with mammalian auditory nerve fibers (Barayeu et al., 2023; Joris et al., 2004). Thus, we expect weakly nonlinear responses for near-threshold stimulation in auditory nerve fibers as well. These could boost or distort responses to two simultaneously presented tones and thus might play a role in forming perception of music.

## Acknowledgements

We thank Tim Hladnik, Henriette Walz, Franziska Kuempfbeck, Fabian Sinz, Laura Seidler, Eva Vennemann, and Ibrahim Tunc for the data they recorded over the years in our lab.

## Founding sources

Supported by SPP 2205 “Evolutionary optimisation of neuronal processing” by the DFG, project number 430157666.

